# Identification of β-III-spectrin actin-binding modulators for treatment of spinocerebellar ataxia

**DOI:** 10.1101/2022.11.08.515660

**Authors:** Piyali Guhathakurta, Robyn T. Rebbeck, Sarah A. Denha, Amanda R. Keller, Anna L. Carter, Alexandra E. Atang, Bengt Svensson, David D. Thomas, Thomas S. Hays, Adam W. Avery

## Abstract

β-III-spectrin is a key cytoskeletal protein that localizes to the soma and dendrites of cerebellar Purkinje cells, and is required for dendritic arborization and signaling. A spinocerebellar ataxia type 5 (SCA5) L253P mutation in the cytoskeletal protein β-III-spectrin causes high-affinity actin binding. Previously we reported a cell-based fluorescence assay for identification of small molecule actin-binding modulators of the L253P mutant β-III-spectrin. Here we describe a complementary, *in vitro*, fluorescence resonance energy transfer (FRET) assay that uses purified L253P β-III-spectrin actin-binding domain (ABD) and F-actin. To validate the assay, we screened a 2,684-compound library of FDA-approved drugs. Importantly, the screening identified numerous compounds that decreased FRET between fluorescently labeled L253P ABD and F-actin. The activity and target of multiple Hit compounds were confirmed in orthologous co-sedimentation actin-binding assays. Through future medicinal chemistry, the Hit compounds can potentially be developed into a SCA5-specific therapeutic. Furthermore, our validated FRET-based *in vitro* HTS platform is poised for screening large compound libraries for β-III-spectrin ABD modulators.

## Introduction

β-III-spectrin is a key actin-crosslinking protein that localizes to the dendrites and soma of cerebellar Purkinje cells (1). Mouse models showed that β-III-spectrin is required for Purkinje cell dendritic arborization and proper postsynaptic localization of multiple membrane proteins (2,3). Autosomal dominant mutations is *SPTBN2* gene encoding β-III-spectrin cause the neurodegenerative disease, spinocerebellar ataxia type 5 (SCA5) (4). SCA5 causes degeneration of Purkinje cells and an associated, progressive limb and gait ataxia. Currently there is no cure or therapy for SCA5.

SCA5-associated mutations localize to the β-III-spectrin N-terminal actin-binding domain (ABD) and spectrin-repeat domains. We previously showed that the ABD-localized L253P mutation causes high-affinity actin binding (5). L253P is positioned at the interface of the two calponin homology domains (CH1 and CH2) comprising the ABD. Our biochemical and biophysical studies showed that L253P causes high-affinity actin binding by opening the CH1-CH2 interface, allowing CH1 to directly bind actin (6). Numerous additional SCA5 mutations localize to the CH1-CH2 interface, suggesting that high-affinity actin binding may be a shared molecular consequence of the ABD-localized SCA5 mutations (7). Thus, it is likely that a small-molecule compound that reduces binding of β-III-spectrin L253P mutant to actin will be effective as a SCA5 therapeutic for other ABD mutants.

We previously reported the development of a cell-based high-throughput screening (HTS) platform to identify small-molecule actin-binding modulators of the L253P ABD (8). In this assay, HEK293 cells are transiently transfected with GFP-ABD (donor) and an actin binding peptide, Lifeact, that is N-terminally labeled with mCherry (acceptor). Binding of GFP-ABD and Lifeact-mCherry on neighboring actin protomers resulted in ∼12% FRET efficiency. Specificity of the FRET signal was demonstrated by the actin-severing compound swinholide A, which greatly reduced the FRET signal. Screening of a 1280-compound library identified Hits that reduced Lifeact binding to actin, and that the assay is sensitive to compounds that cause cell lysis (8).

Here we report the development of a complementary and more sensitive FRET-based HTS screening platform that uses purified L253P ABD and F-actin. Key features include that the assay 1) has greatly increased FRET efficiency, 2) consists of only two components (fluorescently labeled ABD and actin), and 3) is insensitive to compound cytotoxicity and poor cell permeability. Screening of a library of FDA-approved compounds and further evaluation with co-sedimentation assays established the validity of the current assay for detection of small molecules that reduce binding of L253P ABD to actin.

## Results

### Biosensor validation for high-throughput compatibility by swinholide A

To complement our live cell ABD biosensor, we developed an *in vitro* FRET based biosensor using purified ABD and F-actin. A FRET donor construct consisting of the green fluorescent protein, mNeonGreen (mNG), fused to the N-terminus of the mutant ABD (mNG-ABD-L253P), was incubated with the phalloidin-stabilized filamentous actin (F-actin). Actin was labelled with an acceptor dye Alexa Flour 568 (AF568) at residue C374. With mNG-ABD-L253P bound to F-actin, we predicted that each of the mNG fluorescent protein (donor) would be within 7-9 nm distance of AF568 (acceptor) on actin-C374, on the same actin monomer and the neighbouring actin monomers (Figure 1). mNG-ABD-L253P was excited with a 473 nm laser and FRET was measured as a reduction in the fluorescence lifetime (FLT) of the donor (τ_D_), in the presence of the acceptor, AF568-actin. With multiple FRET acceptors within the range of the FRET donor, this aligned with the high level (54%) of FRET observed following 20 min of incubation between 0.5 μM mNG-ABD-L253P and 1 μM AF568-actin (Figure 2A).

**Figure 1.**
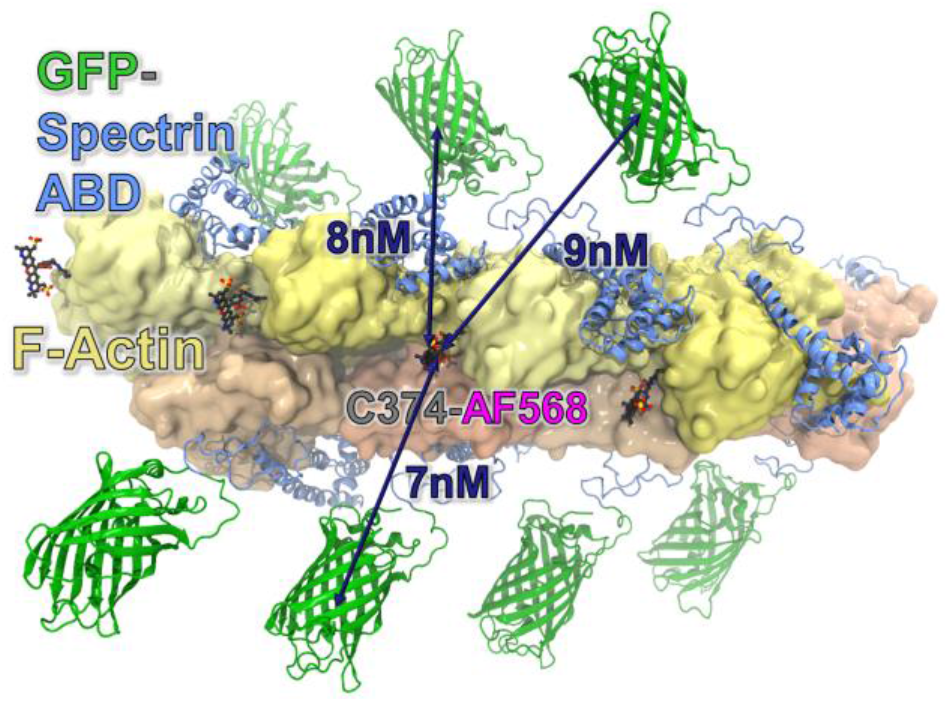
Model of *in vitro* ABD biosensor. Model of mNeonGreen(GFP)-ABD-L253P bound to F-actin labeled with AF568 at C374. Position of β-III-spectrin ABD L253P on actin was based on the cryo-EM structures 6ANU (6), and modeled using DS Visualizer (Dassault Systemes, San Diego, CA, USA). Molecular figure was generated using VMD (35).

**Figure 2.**
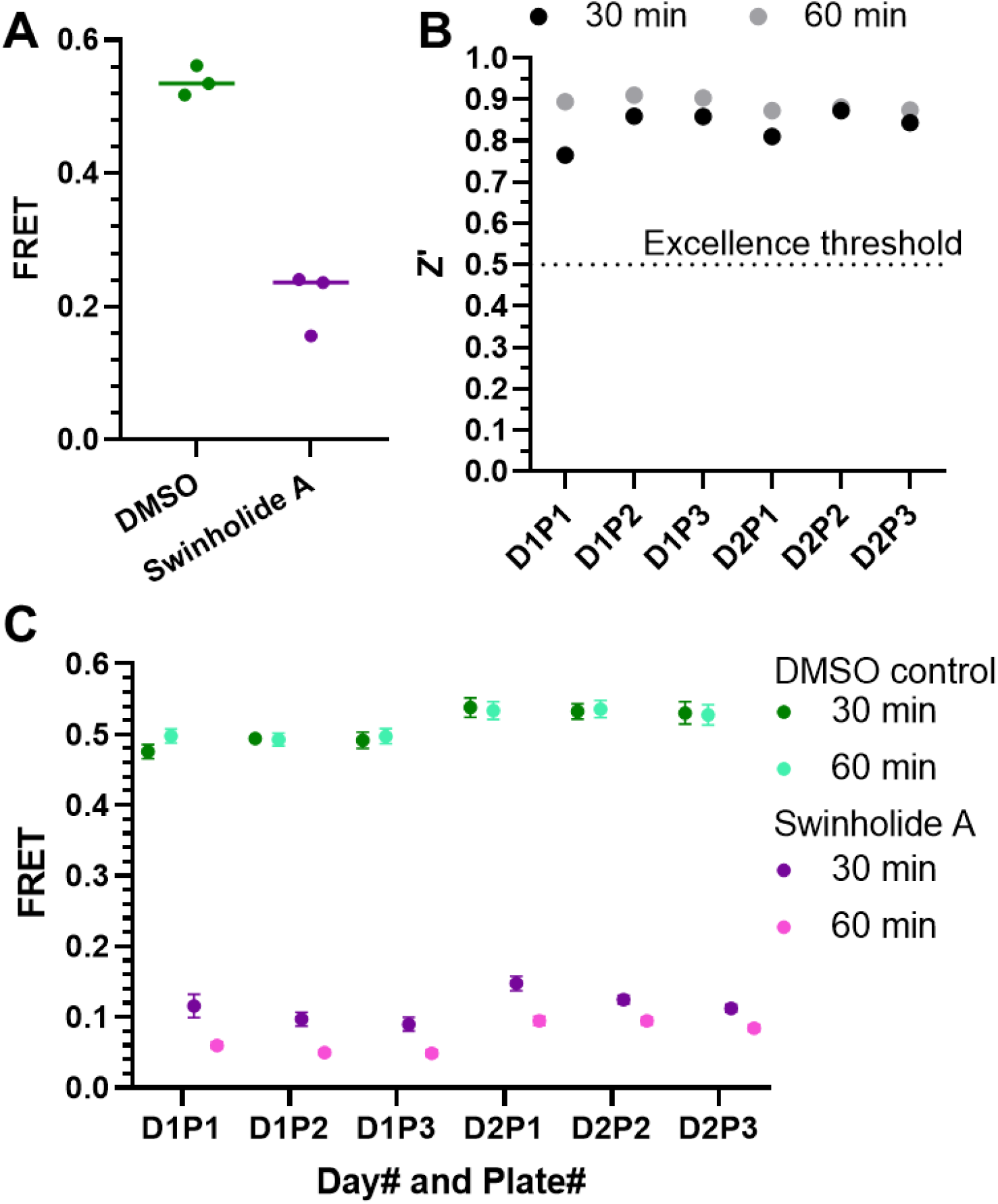
Swinholide A is a reproducible, positive control tool compound for FRET assays in 1536 well plates. **A**. FRET between mNG-ABD-L253P and AF568-actin is reduced by swinholide A. For each plate, each condition was loaded over 27 wells on a 1536 well plate, n=3 plates. **B and C**. Plot of Z’-factor (B) and FRET (C) value per 1536-well plate over two days for 5 μM swinholide A loaded over 32 wells vs DMSO control loaded over 224 wells. FLT were acquired at 30 and 60 min time point after addition of swinholide A or DMSO control.

To test the compatibility of the mNG-ABD-L253P and AF568-actin FRET assay for the primary compound screening, we tested the response of our biosensor to the F-actin severing compound, swinholide A. This compound was previously established as a tool compound for our live cell FRET assay (8). As shown in Figure 2A, 20 min incubation with swinholide A reduced the FRET signal by 60.0±10.2%. Thus, swinholide A was used, here, as a tool compound in the current assay for further characterization.

To gauge HTS assay robustness, we used swinholide A and the FRET biosensor in 1536-well plates to measure the Zʹ value, which factors the signal window and data variation between control and tool compound effect. Classically, a value of 0.5 ≤ Z′ < 1 indicates an excellent assay that is ready for large-scale HTS (9). For evaluating reproducibility, we acquired the FLT values, at 30 and 60 min post loading, on three plates per day for two days. As shown in Figure 2B, the Z’ per plate ranged from 0.77-0.91, easily exceeding the 0.5 Z’-factor excellence threshold. Reading the FLT values at 60 min instead of 30 min post plate loading marginally increased the Z’ value by 6.7% (p = 0.022). This is reflected by the slightly larger effect of swinholide A at the longer time point (Figure 2B). Importantly, FRET values were highly reproducible when measured in repeat tests performed on the same day, or on different days, using different preparations of AF568-actin (Figure 2C). Overall, these results indicate the compatibility of the assay for HTS.

### HTS performance

To test the performance of the FRET assay in HTS, we screened the 2684 compound Selleck FDA and clinical drug library for modulators that alter actin binding of β-III spectrin ABD L253P mutant. This library is desirable for pilot screening, as the library compounds all have a rich research history, and may directly lead into further studies into therapeutic potential for SCA5. The compounds, together with DMSO controls, were dispensed in 15 nL volumes into individual wells of three 1536-well microplates and stored at -20°C until use. Following plate thaw, 5 μL of mNG-ABD L253P alone (Donor only), or mNG-ABD L253P and AF568-actin (Donor + Acceptor) assay mix were loaded into each well via a Multi-drop liquid dispenser. Final compound concentration equalled 30 µM. Plate loading times were staggered to allow for 6-min FLT acquisition time between plates. A time course of compound effects (at 30 and 60 min post load) on FLT was acquired using a high-speed, high-precision fluorescence lifetime plate reader (FLT-PR). This technology has been advanced to high density 1536-well plates in recent years for successful HTS using a range of protein biosensors including β-III spectrin ABD in mammalian cells (8), sarcoplasmic reticulum Ca-ATPase (10-13), ryanodine receptor (9,14), actin (15), cardiac myosin-binding protein C (16), tumor necrosis factor receptor 1 (17,18), and tau (19).

Interfering fluorescent compounds were identified as compounds that altered the fluorescence spectrum by > 3SD, as previously described (9-11,14,20). The effects of the compounds on the FRET-biosensor are shown in Figure 3. For most compounds, there was little variation in the magnitude of the FRET-effect at the 30 or 60 min incubation time (Figure 3A). This is also reflected by the similar number of Hits identified at either time point (Table 1).

**Table 1.** Number (#) of Hits and Hit reproducibility for 3, 5 and 7 standard deviation (SD) thresholds

| **Standard Deviation** |  | **SELLECK Run 1** | **SELLECK Run 2** |
| --- | --- | --- | --- |
| 3SD | # of Hits 30 min (hit rate%) | 492 (18.6%) | 498 (18.8%) |
|  | # of Hits 60 min (hit rate%) | 523 (19.8%) | 501 (18.9%) |
|  | % of Repeated Hits in 2 runs – 30 min | 45.3% | 44.8% |
|  | % of Repeated Hits in 2 runs – 60 min | 49.5% | 51.7% |
| 5SD | # of Hits 30 min (hit rate%) | 247 (10.3%) | 248 (9.4%) |
|  | # of Hits 60 min (hit rate%) | 288 (10.9%) | 250 (9.5%) |
|  | % of Repeated Hits in 2 runs – 30 min | 38.5% | 38.0% |
|  | % of Repeated Hits in 2 runs – 60 min | 38.2% | 44.0% |
| 7SD | # of Hits 30 min (hit rate%) | 164 (6.3%) | 155 (5.8%) |
|  | # of Hits 60 min (hit rate%) | 155 (5.9%) | 145 (5.5%) |
|  | % of Repeated Hits in 2 runs – 30 min | 29.3% | 31.0% |
|  | % of Repeated Hits in 2 runs – 60 min | 36.8% | 39.3% |

**Figure 3.**
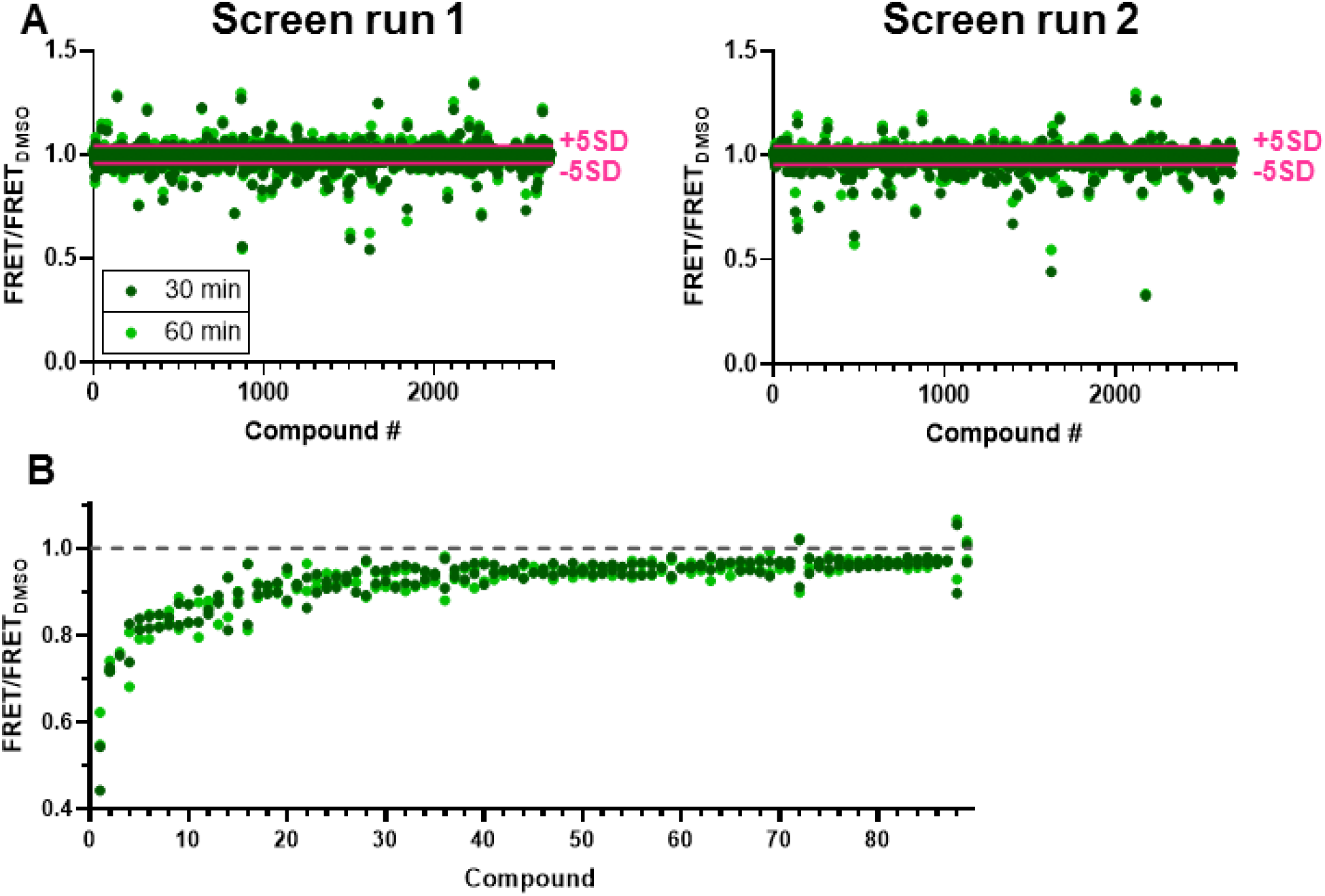
HTS performance validation of *in vitro* ABD biosensor FRET assay using the Selleck Clinical and FDA-approved Drug library in 1536-well plates. FLT data was acquired at two time points following FRET assay loading into 1536-well plates that were pre-loaded with 2684-compound library (30 μM final) or DMSO control. **A**. FRET response to Selleck compounds, with interfering compounds removed, for run 1 (left panel) and run 2 (right panel). FRET response beyond the 5 standard deviation (SD) threshold (pink line) demonstrates that most Hit compounds have little time dependent effect. **B**. Relative FRET effect of Selleck Hits that were identified (with 5SD threshold) as decreasing FRET in both screen runs, with an additional 17 compounds added due to potent effect (> 7 SD) in one screen. Data shown as relative to DMSO control (gray dotted line). Chemical Structures shown in Figures S3-11.

To cast a wide net for possible modulators, we adopted a 5SD threshold for reproducible Hits. Notably, the resulting number of Hits (135 compounds) is higher than the optimal Hit rate (<3%) (21). However, this is likely the result of having a high FRET signal window (∼54%) relative to previous biosensors (<20%) (8,9,14). An additional contributing factor is that we did not remove compounds that alter donor-only FLT, in order to include potential compounds that impact donor-only FLT in the process of binding and structurally modifying the ABD.

In addition to the reproducible compounds, we focused on 17 compounds that potently (> 7SD) altered FRET in one of the two screens. Thus, prioritized compounds were selected from a list of 152 Hit compounds from the screens. Considering the average of the compound effects, this resulted in 89 FRET decreasers and 63 FRET increasers (Figure 3B, Figures S3-16).

An additional factor that may contribute to the higher Hit percentage is that this library contains different salt forms for several compounds and several groups of similar chemical types. Indeed, we identified several salt forms and analogues of chemical types or drug classes. Because our ultimate interest is to identify compounds that reduce the affinity of mutant β-III-spectrin for actin, we further characterized the Hits that reduced FRET.

### FRET dose-response assay

38 Hits that showed the most potent response for their salt form or drug class on FRET were selected for purchase and further validation by first acquiring the FRET response to a range of Hit compound concentrations (0.01-100 µM) under the same assay conditions as used in the primary screen. As shown in Figure 4, 16 of the repurchased Hits reduced FRET by >20%, notably with similar effects as observed in the primary screens. Of the remaining 22 compounds, 19 compounds significantly reduced FRET (Figure S17). Encouragingly, 92% of Hits were verified upon repurchase.

**Figure 4.**
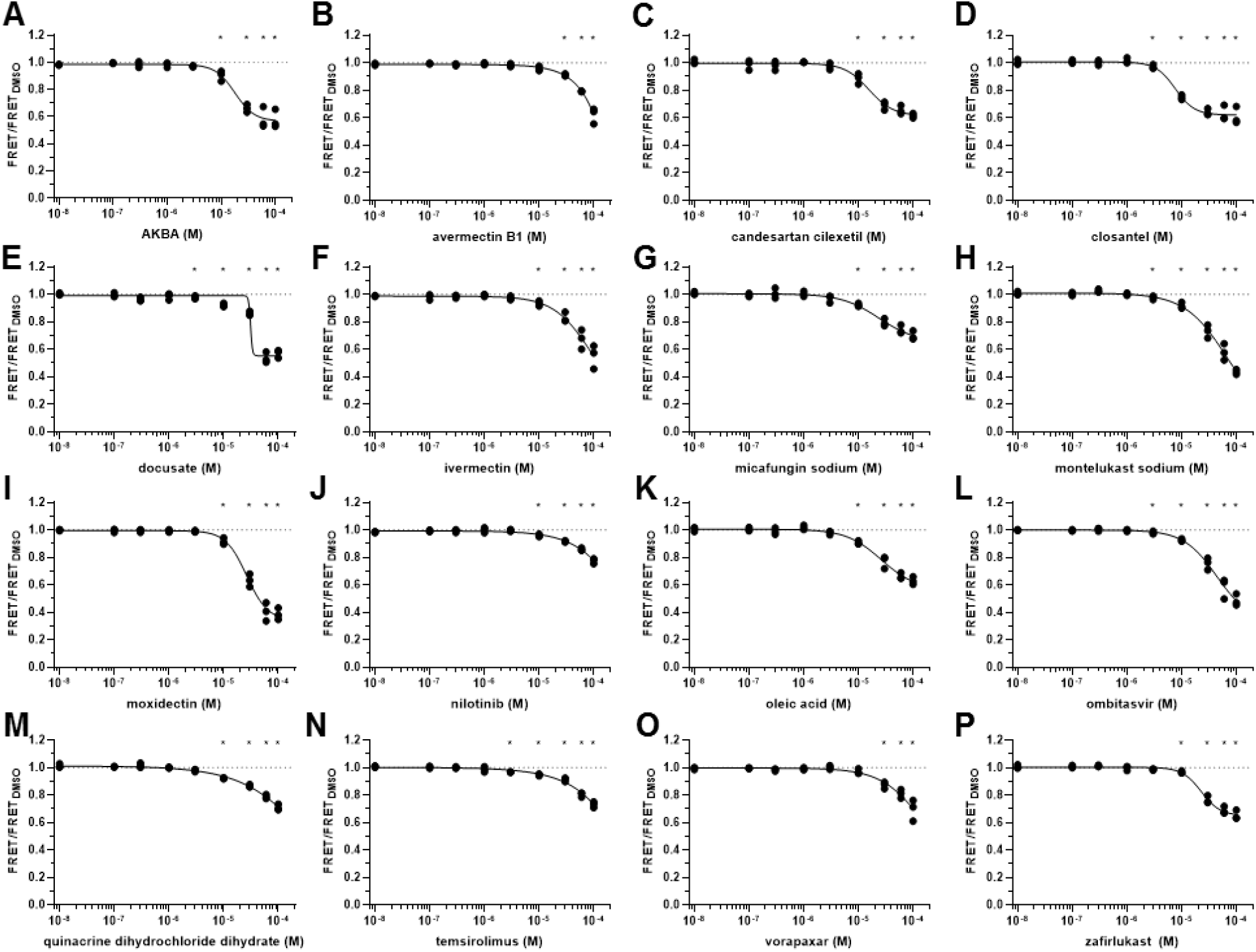
FRET dose response of Hit compounds that decrease FRET by greater than 20%. **A-P**. Dose response of Hit compounds AKBA (**A**), avermectin B1 (**B**), candesartan (**C**), closantel (**D**), docusate (**E**), ivermectin (**F**), micafungin (**G**), montelukast (**H**), moxidectin (**I**), nilotinib (**J**), oleic acid (**K**), ombitasvir (**L**), quinacrine (**M**), temsirolimus (**N**), vorapaxar (**O**), and zafirlukast (**P**) were tested on the *in vitro* ABD biosensor FRET. The chemical structures of these Hit compounds are shown in Figure S3-16. *p< 0.05 relative to DMSO control, using Student’s unpaired T-test. Data shown as individual data points, n=3.

### Confirmation of Hit compound activity in orthologous binding assay

To confirm the activity of Hits to reduce binding of the L253P ABD to F-actin, co-sedimentation assays were performed using purified ABD and F-actin (Figure 5). 2 µM ABD was incubated with 1.2 µM F-actin in the presence of 100 µM compound or DMSO. Most of the 33 compounds that reduced binding in the FRET assay either decreased or increased co-sedimentation of the mutant ABD with F-actin.

**Figure 5.**
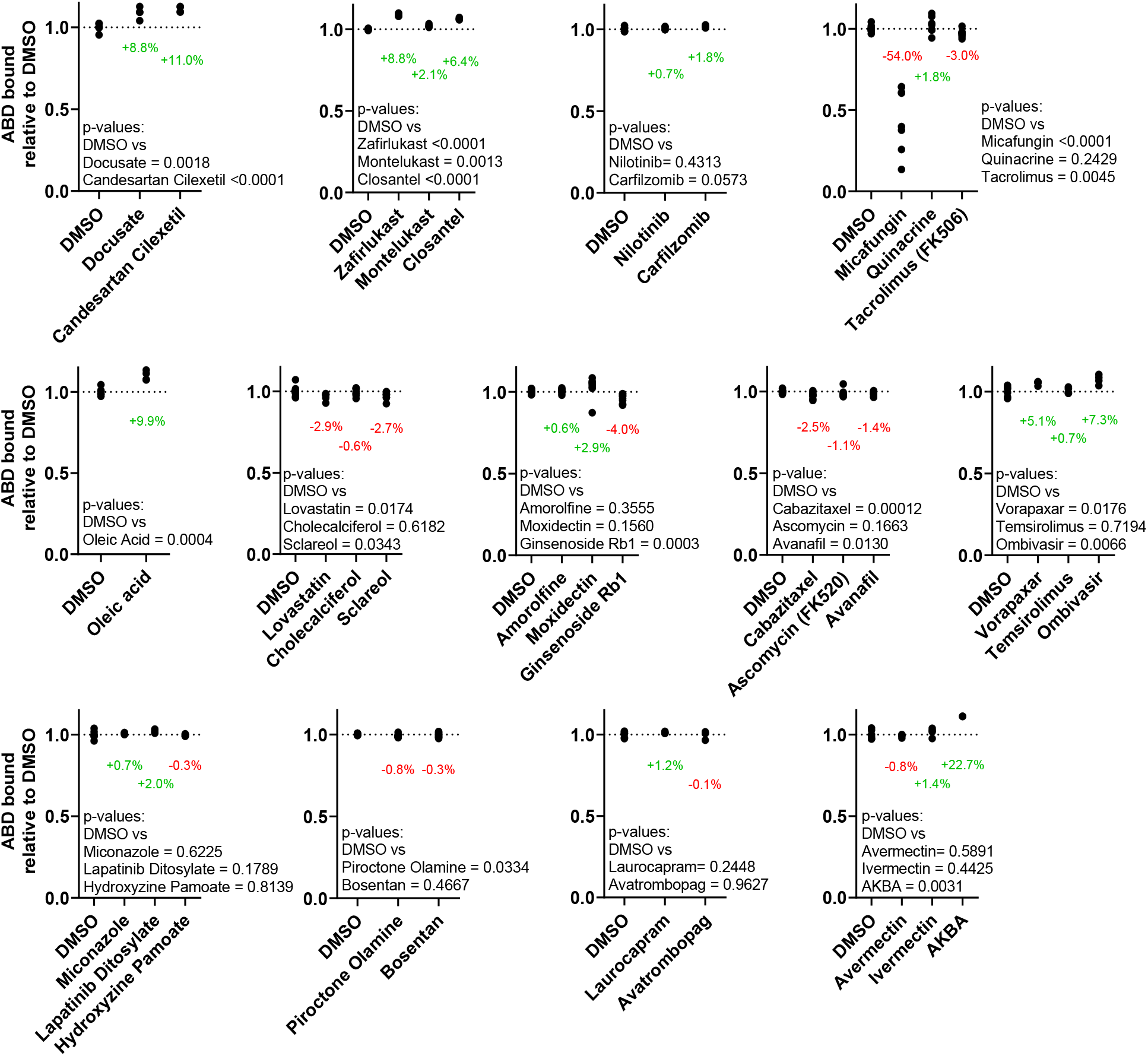
Confirmation of Hit compounds in actin co-sedimentation assays. Co-sedimentation of L253P ABD and actin shows that most Hit compounds (100 μM) decrease or increase co-sedimentation of the L253P ABD (2 μM) with F-actin (1.2 μM). Data is shown as relative to DMSO, n= 3-12.

Eleven compounds caused a significant (2-22%) increase in co-sedimentation of the ABD with actin. This increase in ABD co-sedimentation either reflects an effect of the compounds to increase binding of the ABD to F-actin (counter the FRET results) or compound-induced aggregation of the mutant ABD. To examine the aggregation effects of these compounds, we tested whether the compounds caused ABD sedimentation in the absence of actin (Figure 6). All of the compounds caused the ABD to enter the pellet in the absence of actin, confirming that the Hit compounds were causing ABD aggregation. Consistently, many of the compounds that we experimentally determined to cause ABD aggregation are known or predicted aggregators in the Aggregator Database (22) (Table S1).

**Figure 6.**
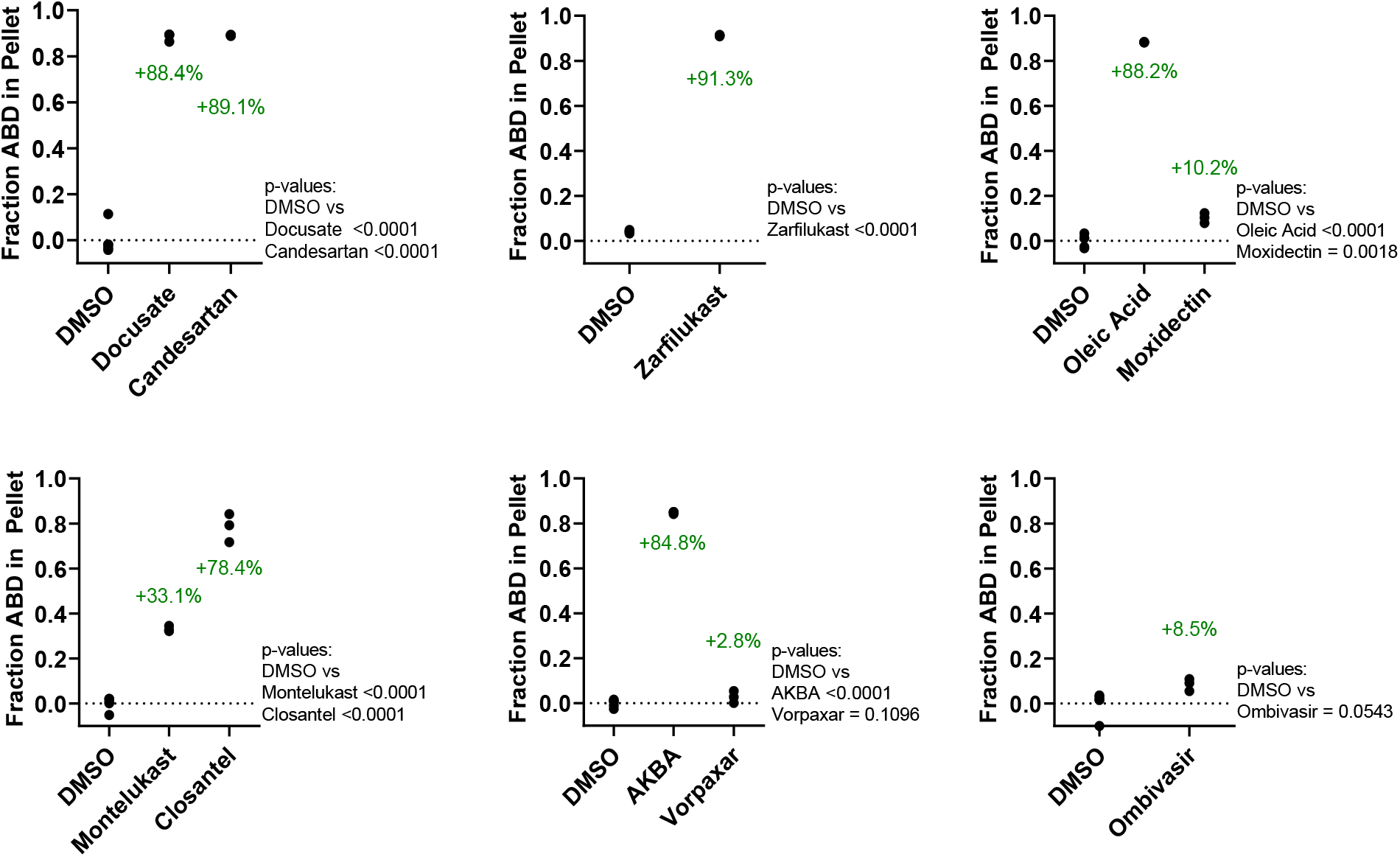
Aggregation assay of Hit compounds. For Hit compounds that caused the L253P ABD to enter pellet in actin co-sedimentation assays, co-sedimentation reactions were repeated in the absence of actin. All tested compounds (100 µM) caused the L253P ABD to enter pellet in the absence of actin. Data is shown as relative to DMSO, n= 3-6.

Nine compounds reduced actin binding in co-sedimentation assays significantly, by ∼1 to ∼50%. Of these compounds, Micafungin had the largest effect, decreasing ABD co-sedimentation by ∼50%. Other compounds that caused a significant reduction in ABD co-sedimentation included Tacrolimus, Lovastatin, Sclareol, Ginsenoside Rb1, Cabazitaxel, Avanafil and Piroctone (Figure 5). Ascomycin also caused average ABD binding to be reduced, although the effect fell short of statistical significance. For these compounds that reduced ABD binding in the co-sedimentation assays, similar effect sizes were observed in the FRET binding assays (Figure 4, Figure 7, Figure S17). Of the nine compounds, Ginsenoside Rb1 had the lowest EC50, ∼3 µM (Figure 7, Figure S17). None of the Hit compounds that reduced ABD co-sedimentation are known or similar to known aggregators in the Aggregator Database (Table S1). These nine Hit compounds are thus of greatest interest as potential modulators of L253P mutant β-III-spectrin actin-binding activity.

**Figure 7.**
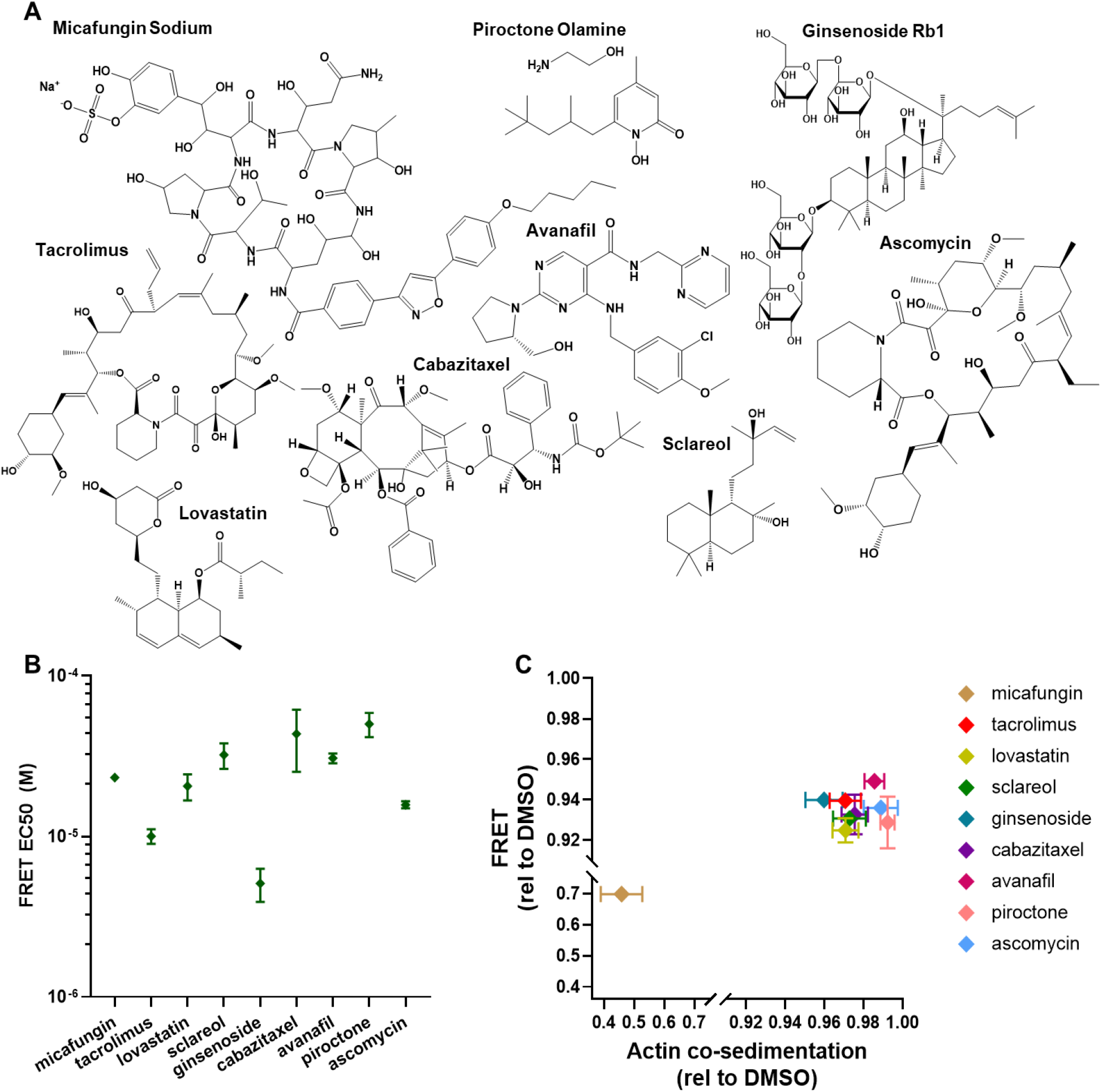
Summary of compounds that reduce ABD-actin binding. **A**. Compound structures. **B**. FRET EC50 from Hill fits of data shown in Figure 4 and Figure S17. Data shown as mean ± SEM, n=3. **C**. Comparison of 100 μM compound on actin co-sedimentation vs FRET, data shown in Figures 4 and 5, and Figure S17.

## Discussion

We have developed a high-precision, FRET-based HTS assay that monitors binding of recombinant mNeonGreen-tagged L253P β-III-spectrin ABD to actin filaments covalently linked to a fluorescent dye Alex Fluor 568. In this assay, binding of the L253P ABD to actin filaments results in a ∼50% FRET efficiency. This FRET signal is abolished by the F-actin severing compound, swinholide A, in agreement with our prior cryo-EM model showing that ABD actin binding involves contact with two adjacent actin subunits within the filament (6). The large FRET signal makes the assay amenable to HTS, as indicated by the high Z’ values achieved when the assay is performed in 1536-well microplates.

We validated the assay for HTS by screening Selleck’s 2684-compound FDA and clinical library. Duplicate screens led to the identification of 9 compounds with confirmed activity to decrease L253P ABD and actin binding. This highlights the effectiveness of this assay at identifying compounds with desirable activity. In addition, these Hit compounds support that the ABD-actin interaction is a druggable target. The successful identification of Hit compounds in this small compound library screen warrants further screening of larger compound libraries to increase the number and structural diversity of the Hit compounds.

Of the nine Hit compounds that reduced L253P ABD and actin binding (Figure 5 and 7), micafungin stands out as having the largest effect size, reducing actin binding by ∼50%. Micafungin is a semi-synthetic compound approved by the FDA as an antifungal agent that works by inhibiting synthesis of the fungal cell wall component, 1,3-β-D-glucan (23). Micafungin is intravenously injected to treat systemic fungal infections, and is reported to be tolerated at high dose with little adverse side effect (24). Micafungin’s large size (1292.26 Da) probably limits its ability to cross the blood brain barrier. However, Micafungin can be detected at low level in the CNS (25).

Ginsenoside Rb1 stood out as the Hit with the lowest EC50, ∼3 µM, and reduced actin binding by 4-6% in actin co-sedimentation and FRET assays. Ginsenosides are bioactive components of the ginseng plant. Intriguingly, numerous studies in mammalian cell systems suggest that Ginsenoside Rb1 has neuroprotective properties (26). For example, Ginsenoside Rb1 was reported to ameliorate motor deficits in a mouse model of Parkinson’s disease, potentially by reducing glutamate-mediated neurotoxicity by upregulated expression of the GLT-1 glutamate transporter (27). Purkinje cell loss in SCA5 pathogenesis has also been linked to glutamate-mediated neurotoxicity (28,29). It is appealing to hypothesize that Ginsenoside Rb1 could work as a SCA5 therapeutic by both reducing β-III-spectrin actin binding and increasing clearance of glutamate.

Currently, we are in the early phase of SCA5 therapeutic discovery. While the therapeutic potential of current Hit compounds is supported by our *in vitro* assays, we emphasize that none of Hit compounds have been evaluated for efficacy or safety in treating neurodegeneration or ataxia associated with SCA5, in any system. For our current Hit compounds, medicinal chemistry may be necessary to increase compound potency (reduced EC50 and/or increased effect size). Moreover, the activity of the compounds should be tested in cell models that have revealed mislocalization of the L253P mutant β-III-spectrin (30,31). To assess Hit compound therapeutic potential, it is now essential that a SCA5 L253P mouse model be generated. Further, screening of additional, larger compound libraries would increase the number and structural diversity of Hits, ensuring successful development of a SCA5 therapeutic.

### Experimental procedures

#### Expression and purification of mNeonGreen and mNeonGrenn-ABD proteins

The mNeonGreen coding sequence was synthesized at Integrated DNA Technologies and matches Genbank sequence KC295282.1. mNeonGreen was digested with KpnI and EcoRI and subcloned into pcDNA3.1-GFP-ABD L253P (32), after digestion with KpnI and EcoRI to remove GFP. mNeonGreen (mNG) and mNeonGreen-ABD L253P (mNG-ABD L253P) coding sequences were PCR amplified with the forward primer AAACACCTGCAAAAAGGTATGGTGAGCAAGGGCGAGGAGG, and the reverse primer AAATCTAGACTACTTGTACAGCTCGTCCAT GCCC for mNG, or the reverse primer AAATCTAGACTACTTCATCTTGGAGAAGTAATGGTAGTAAG for mNG-ABD L253P. Following digest with Aarl and Xbal, PCR products were ligated into BsaI-digested pE-SUMOpro (LifeSensors). Sequence-verified constructs were transformed into Rosetta 2 (DE3) *E. coli* (Novagen). mNG and mNG-ABD L253P protein expression and purification steps, including removal of SUMO tag, were performed as previously described (32). A final step of buffer exchange was performed for mNG by dialysis in buffer containing 10 mM Tris, pH 7.5, 150 mM NaCl, 2 mM MgCl_2_, and 1 mM DTT using Slide-A-Lyzer, 10,000 MWCO cassettes (Thermo Scientific). For mNG-ABD L253P a final gel filtration step was performed using a S-100 sephacryl column equilibrated with buffer containing 10 mM Tris, pH 7.5, 150 mM NaCl, 2 mM MgCl_2_, and 1 mM DTT. Fractions containing mNG-ABD L253P were concentrated using a Centrifugal Filter Unit, 10,000 MWCO (Millipore). mNG and mNG-ABD L253P protein concentrations were determined through Bradford assay. Purified proteins were supplemented with sucrose (150 mM final), snap frozen in liquid nitrogen and stored at -80 °C until use.

#### Actin preparation and labeling

Actin was prepared from rabbit skeletal muscle by extracting acetone powder in cold water, as described previously (52). 130 μM Alexa Fluor 568 C5 maleimide (Invitrogen), freshly dissolved in dimethylformamide (DMF), was added to 65 μM F-actin and the sample was incubated for 30 min at 25°C and then 18 h at 4°C. Labeling was terminated by adding 10 mM DTT, and actin was ultracentrifuged for 30 min at 350,000 × g. The F-actin pellet was suspended in G-Mg buffer (5 mM Tris, 0.5 mM ATP, 0.2 mM MgCl_2_, pH 7.5) followed by clarification at 300,000 × g for 10 min. Actin was again polymerized for 45 min at 25°C in the presence of 3 mM MgCl_2_ and ultracentrifuged at 350,000 × g for 30 min. F-actin pellet was suspended in F-Mg buffer (3 mM MgCl_2_, 10 mM Tris, pH 7.5) containing 0.2 mM ATP. The labeled F-actin was immediately stabilized against depolymerization and denaturation by adding equimolar phalloidin.

#### Fluorescence data acquisition

Fluorescence lifetime measurements were carried out by a high-precision FLTPR (provided by Photonic Pharma LLC, Minneapolis, MN) (11,33). Donor sample (mNG-β-III-spectrin ABD L253P) was excited with a 473-nm microchip laser (Bright Solutions, Cura Carpignano, Italy), and emission was filtered with 488-nm long pass and 517/20-nm band pass filters (Semrock, Rochester, NY). This instrument enables high-throughput fluorescence lifetime detection at high precision by utilizing a unique direct waveform recording technology (11). The performance of this FLTPR has been previously demonstrated with FRET-based HTS that targets several muscle and non-muscle proteins (8,11,14). In the present study, modifications were made in the instrument to permit 2-channel detection, for the purpose of flagging false Hits due to interference from fluorescent compounds.

#### Screen with SELLECK library

The 2684 SELLECK compounds were received in 96-well plates and reformatted into 1536-well flat, black-bottom polypropylene plates (Greiner Bio-One). In total, 50 nl of each compound solution was dispensed in DMSO using an automated Echo 550 acoustic liquid dispenser (Labcyte). Compounds were formatted into the assay plates, at a final concentration of 30 μM, with the first two and last two columns loaded with DMSO only (compound-free controls). These assay plates were then heat-sealed using a PlateLoc Thermal Microplate Sealer (Agilent Technologies) and stored at –20⁰C. Before screening, compound plates were equilibrated to room temperature (25⁰C). In total, 0.5 μM mNG-ABD1 without or with 1 μM Alexa-568-labeled actin was dispensed by a Multidrop Combi Reagent Dispenser (Thermo Fisher Scientific) into the 1536-well assay plates containing the compounds. Plates were incubated at room temperature for 60 min before recording the data with the FLTPR. A control measurement with 0.5 µM mNG (fluorescent protein only) was performed to eliminate the compounds that affected the environment of the fluorescence protein.

#### HTS data analysis

Waveforms for each well in HTS were convolved with the instrument response function and were fitted by a one-exponential decay function using least-squares minimization (34). The FRET efficiency (*E*) was determined as the fractional decrease of donor fluorescence lifetime (τ_D_), due to the presence of acceptor fluorophore (τ_DA_):

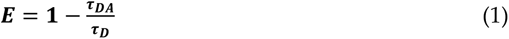

Assay quality was determined based on FRET assay samples in wells pre-loaded with control (DMSO) and tested tool compound, as indexed by the Z΄ factor:

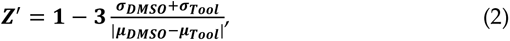

where σ_DMSO_ and σ_Tool_ are the SDs of the DMSO τ_DA_ and tool compound τ_DA_, respectively; µ_DMSO_ and µ_Tool_ are the means of the DMSO τ_DA_ and tool compound τ_DA_, respectively. A compound was considered a Hit if it changed τ_DA_ by > 5 SD relative to that of control τ_DA_ that were exposed to 0.3% DMSO.

#### Compound’s concentration–response assay

The Hit compounds were purchased and dissolved in DMSO to make a 10 mM stock solution, which was serially diluted in 96-well mother plates. Hits were screened at nine concentrations (0.01–100 μM), with DMSO controls also loaded. Compounds (1 μl) were transferred from the mother plates into 1536-well plates using a Mosquito HV liquid handler (TTP Labtech Ltd). The same procedure of dispensing as for the pilot screening was applied in the TR-FRET concentration–response assays. The FRET efficiency E was determined as the fractional decrease in donor fluorescence lifetime as described above. Concentration dependence of the FRET (relative to DMSO control) change was fitted using the Hill equation.

#### Co-sedimentation

F-actin co-sedimentation assays were performed as described previously (32) with few modifications. After measuring the concentrations of polymerized F-actin and clarified ABD proteins via Bradford assay, the binding reactions were set-up by combining 2 µM ABD protein, 1.2 µM F-actin, and 100 µM compound or DMSO, to a 60 µL total reaction volume in F-buffer (10 mM Tris, pH 7.5, 150 mM NaCl, 0.5 mM ATP, 2 mM MgCl_2_, and 1 mM DTT). The binding reactions were incubated at 21⁰C for 30 min, then centrifuged at 100,000 x g at 25°C to pellet F-actin. The supernatants containing the unbound ABD were collected and mixed with 4X Laemmli sample buffer. The unbound ABD samples were separated by SDS-PAGE and stained with Coomassie blue R-250 stain for 2 hours. Protein band intensities were measured using Image Studio Lite version 5.2 software after imaging the gels using the 680 nm channel on Azure Sapphire scanner. Fraction ABD bound was measured after converting fluorescence intensity measurements to concentrations using standard curve generated from a Coomassie blue stained gel containing varying amounts ABD protein in the absence of actin or compound. The data were plotted in Prism 9 (GraphPad) relative to DMSO control average. The aggregation assays were performed as the co-sedimentation assays but without F-actin.

### Analysis and presentation of data

Data is presented as mean ± SD or ± SEM, as indicated. For statistical difference determination, unpaired Student’s T-test was performed. Statistical analyses were performed with GraphPad Prism and Origin. Significance was accepted at P < 0.05. EC_50_ values were derived from fits to Hill equations.

## Data availability

All data are contained within the manuscript.

## Supporting Information

This article contains supporting information

## Acknowledgments

We are grateful to Samantha Yuen and Jonathan Solberg for preparing compound assay plates.

## Author contributions

RTR, PG and AWA conceived and designed the experiments. PG, ALC, ARK, SAD, AEA, RTR and AWA acquired data and interpreted results. BS generated the mNeonGreen-ABD-L253P and F-actin Alexa Fluor 568 model. RTR and AWA prepared the manuscript draft. PG, DDT and TSH reviewed and edited the manuscript. All authors have read and agreed to the published version of the manuscript.

## Funding and additional information

This work was supported by NIH grants R61NS111075, R33NS111075 and R01GM044757 to TSH, R01HL139065 (formerly GM027906) and R37AG026160 to DDT, and R15NS116511 to AWA.

## Abbreviations and nomenclature

ABD: actin-binding domain
CH: calponin homology
DMSO: dimethyl sulfoxide
F-actin: actin filaments
FLT: fluorescent lifetime
FRET: fluorescence resonance energy transfer
HTS: high-throughput screening
PR: plate reader
SCA5: spinocerebellar ataxia type 5

