## Supplemental Data for "Identification of β-III-spectrin actin-binding modulators for treatment of spinocerebellar ataxia"


**Running title**: ***In vitro*** **drug discovery targeting β-III-spectrin**

**Table S1. Aggregator Database results for Hit compounds.** Many of the Hits that we determined experimentally to be ABD aggregators are either known aggregators or somewhat/very similar to known aggregators. In contrast, Hits that did not cause ABD aggregation are not similar to known aggregators. High logP indicates compound is hydrophobic and has increased potential to act as aggregator.

| Hit | Known Aggregator? |
|---|---|
| Candesartan | Very similar to known aggregator |
| Oleic acid | Very similar to known aggregator |
| Docusate | Not similar to known aggregator, but high logP |
| Zafirlukast | Very similar to known aggregator |
| Montelukast | Not similar to known aggregator, but high logP |
| Closantel | Data not available |
| Moxidectin | Not similar to known aggregator, but high logP |
| Micafungin | Not similar to known aggregator |
| Quinacrine | Not similar to known aggregator, but high logP |
| Tacrolimus | Not similar to known aggregator, but high logP |
| Lovastatin | Not similar to known aggregator, but high logP |
| Sclareol | Not similar to known aggregator, but high logP |
| Ginsenoside Rb1 | Not similar to known aggregator |
| Cabazitaxel | Not similar to known aggregator, but high logP |
| Ascomycin | Not similar to known aggregator, but high logP |
| Avanafil | Not similar to known aggregator, but high logP |
| Piroctone | Not similar to known aggregator |
| Bosetan | Very similar to known aggregator |

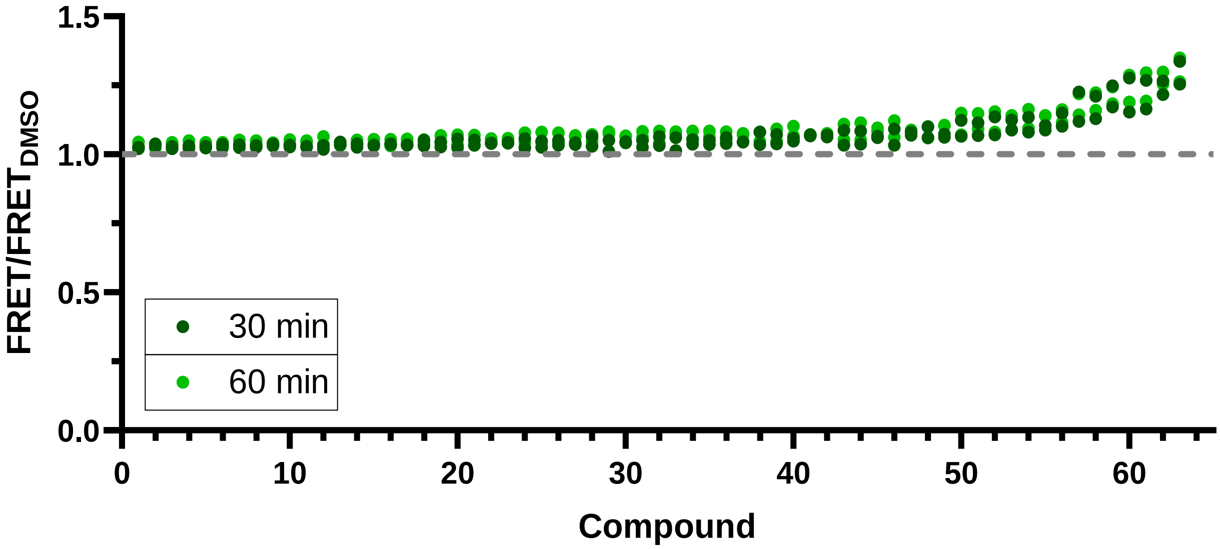


**Figure S1. Hits that reproducibly increased *in vitro* ABD biosensor FRET between two screens of the Selleck library.** Relative FRET effect of Selleck Hits that were identified (with 5SD threshold) increasing FRET in both screen runs. Data is shown as relative to DMSO control (gray dotted line). Chemical Structures shown in Figures S11-16.

| | |
|---|---|
| 1 | Piperonyl butoxide |
| 2 | Dipyridamole |
| 3 | **Candesartan Cilexetil** |
| 4 | **Oleic Acid** |
| 5 | **Micafungin Sodium** |
| 6 | **Docusate Sodium** |
| 7 | **Montelukast Sodium** |
| 8 | Sodium lauryl sulfate |
| 9 | **Quinacrine 2HCl** |
| 10 | Quinacrine 2HCl 2H2O |
| 11 | **Zafirlukast** |
| 12 | Dalbavancin |
| 13 | Erythromycin estolate |
| 14 | Tyrosol |
| 15 | Cefsulodin sodium |
| 16 | Ledipasvir (GS5885) |
| 17 | **Temsirolimus** |
| 18 | Thimerosal |
| 19 | Montelukast |
| 20 | Zotarolimus(ABT-578) |
| 21 | **Ombitasvir (ABT-267)** |
| 22 | **Batyl alcohol** |
| 23 | **cis-Anethole** |
| 24 | Bronopol |
| 25 | Anidulafungin (LY303366) |
| 26 | Simvastatin |
| 27 | Ridaforolimus (Deforolimus) |
| 28 | **Moxidectin** |
| 29 | Aprepitant |
| 30 | Asunaprevir |
| 31 | Rapamycin (Sirolimus) |
| 32 | Simeprevir |
| 33 | Manidipine 2HCl |
| 34 | Clindamycin palmitate HCl |
| 35 | Ertapenem sodium |
| 36 | **Ethotoin** |
| 37 | Hydroxyprogesterone caproate |
| 38 | Latanoprost |
| 39 | Cilnidipine |
| 40 | Ouabain |
| 41 | **Carfilzomib** |
| 42 | Dabigatran Etexilate |
| 43 | Thiostrepton |
| 44 | Rolapitant |
| 45 | Daclatasvir (BMS-790052) |
| 46 | Celecoxib |
| 47 | Tafluprost |
| 48 | **Vorapaxar** |
| 49 | Grazoprevir |
| 50 | Ibuprofen piconol |
| 51 | **Cabazitaxel** |
| 52 | **Ivermectin** |
| 53 | **Avermectin B1(Abamectin)** |
| 54 | **Tacrolimus (FK506)** |
| 55 | **Clonidine HCl** |
| 56 | Troglitazone (CS-045) |
| 57 | **cholecalciferol** |
| 58 | Nefazodone hydrochloride |
| 59 | **Laurocapram** |
| 60 | Cabozantinib malate (XL184) |
| 61 | Ketoconazole |
| 62 | Econazole |
| 63 | **Bosentan Hydrate** |
| 64 | **Avanafil** |
| 65 | **Miconazole Nitrate** |
| 66 | Obeticholic Acid |
| 67 | **Ascomycin (FK520)** |
| 68 | Everolimus (RAD001) |
| 69 | **Ginsenoside Rb1** |
| 70 | **Amorolfine HCl** |
| 71 | Cobicistat (GS-9350) |
| 72 | Luteolin |
| 73 | Ampicillin Trihydrate |
| 74 | **Nilotinib hydrochloride** |
| 75 | **Securinine** |
| 76 | **Sclareol** |
| 77 | Pneumocandin B0 |
| 78 | **AKBA** |
| 79 | Calcitriol |
| 80 | **Hydroxyzine pamoate** |
| 81 | Saikosaponin A |
| 82 | Canagliflozin hemihydrate |
| 83 | Efonidipine |
| 84 | **Avatrombopag** |
| 85 | **Lapatinib Ditosylate** |
| 86 | **Lovastatin** |
| 87 | **Piroctone Olamine** |
| 88 | Ceftibuten dihydrate |
| 89 | **Closantel** |
| 90 | Fursultiamine |
| 91 | Triflusal |
| 92 | Succimer |
| 93 | Bromocriptine Mesylate |
| 94 | Cisatracurium Besylate |
| 95 | Amsacrine hydrochloride |
| 96 | Clomipramine HCl |
| 97 | Primaquine Diphosphate |
| 98 | Salmeterol |
| 99 | Doxycycline Hyclate |
| 100 | Retigabine 2HCl |
| 101 | Fangchinoline |
| 102 | Silymarin |
| 103 | Sulbutiamine |
| 104 | Levosimendan |
| 105 | Olodaterol hydrochloride |
| 106 | Pimavanserin |
| 107 | Paroxetine mesylate |
| 108 | Paroxetine HCl |
| 109 | Nintedanib Ethanesulfonate Salt |
| 110 | Nifedipine |
| 111 | pyrvinium |
| 112 | Benzalkonium chloride |
| 113 | Carvedilol Phosphate |
| 114 | (S)-crizotinib |
| 115 | Flupirtine maleate |
| 116 | Tigecycline |
| 117 | Ilaprazole |
| 118 | Methylcobalamin |
| 119 | CP21R7 (CP21) |
| 120 | Sunitinib Malate |
| 121 | Nebivolol HCl |
| 122 | (−)-Norepinephrine |
| 123 | Crizotinib (PF-02341066) |
| 124 | Omadacycline tosylate |
| 125 | Anlotinib (AL3818) 2HCl |
| 126 | Meclocycline Sulfosalicylate |
| 127 | Trichlormethiazide |
| 128 | Vortioxetine HBr |
| 129 | Benzethonium Chloride |
| 130 | Cyclofenil |
| 131 | Mivacurium chloride |
| 132 | Raloxifene HCl |
| 133 | Tegaserod Maleate |
| 134 | Ethidium bromide |
| 135 | Ethacridine lactate monohydrate |
| 136 | Methylene Blue |
| 137 | Bacitracin Zinc |
| 138 | Phenazine methosulfate |
| 139 | Chlorhexidine 2HCl |
| 140 | Domiphen Bromide |
| 141 | Chlorhexidine |
| 142 | Sanguinarine chloride |
| 143 | Fingolimod (FTY720) HCl |
| 144 | 4-Aminophenol |
| 145 | Zinc Undecylenate |
| 146 | Zinc Pyrithione |
| 147 | Otilonium Bromide |
| 148 | Olanexidine HCl semihydrate |
| 149 | Cetrimonium Bromide (CTAB) |
| 150 | Cetylpyridinium Chloride |
| 151 | Pixantrone Maleate |
| 152 | Octenidine Dihydrochloride |

**Figure S2. Compound names for Hits that altered *in vitro* ABD biosensor FRET**, shown in Figures 3 and S1.Thirty eight Hits that were chosen for further testing are indicated by green text.

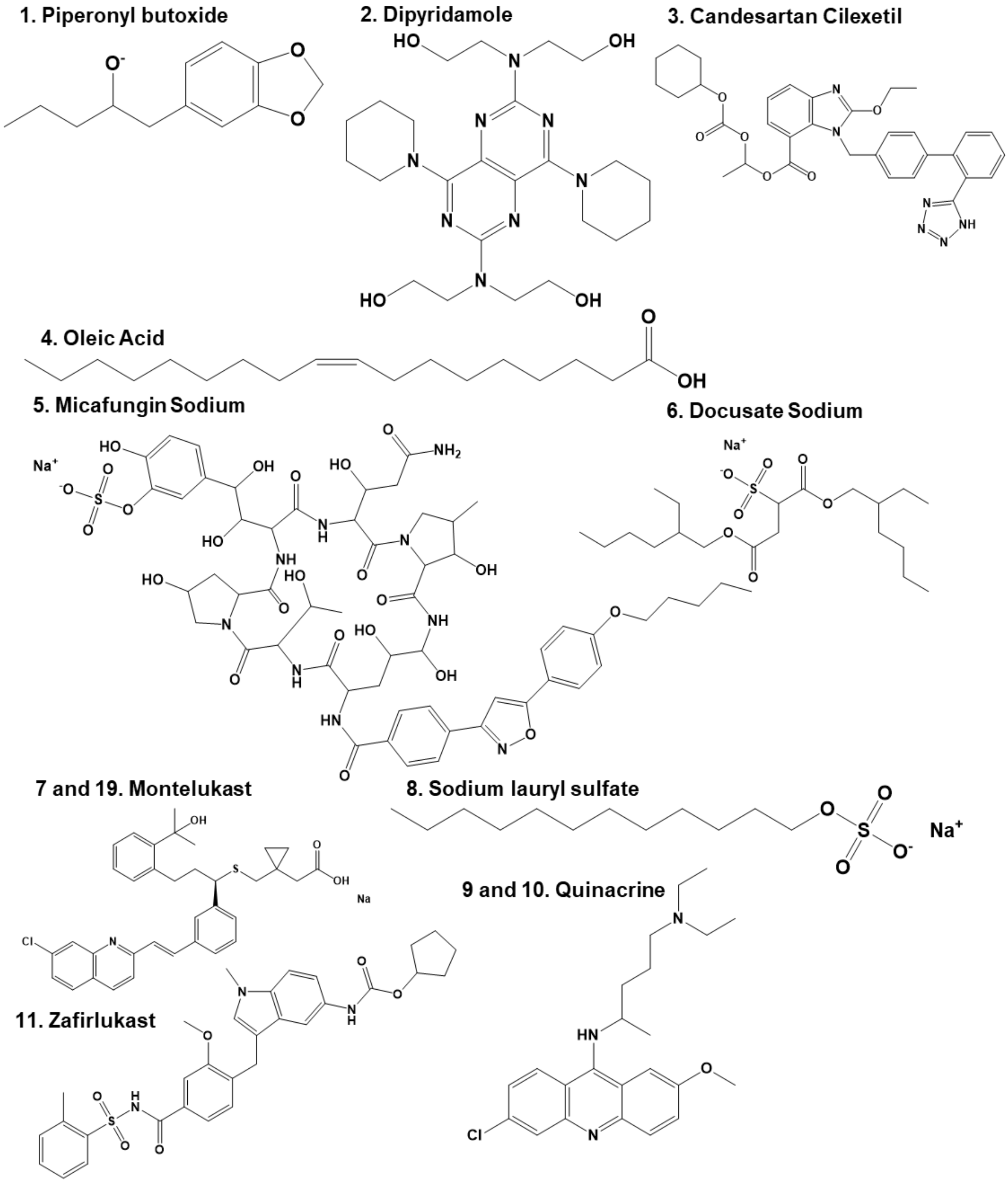


**Figure S3. Chemical structures of Selleck screen Hits.**

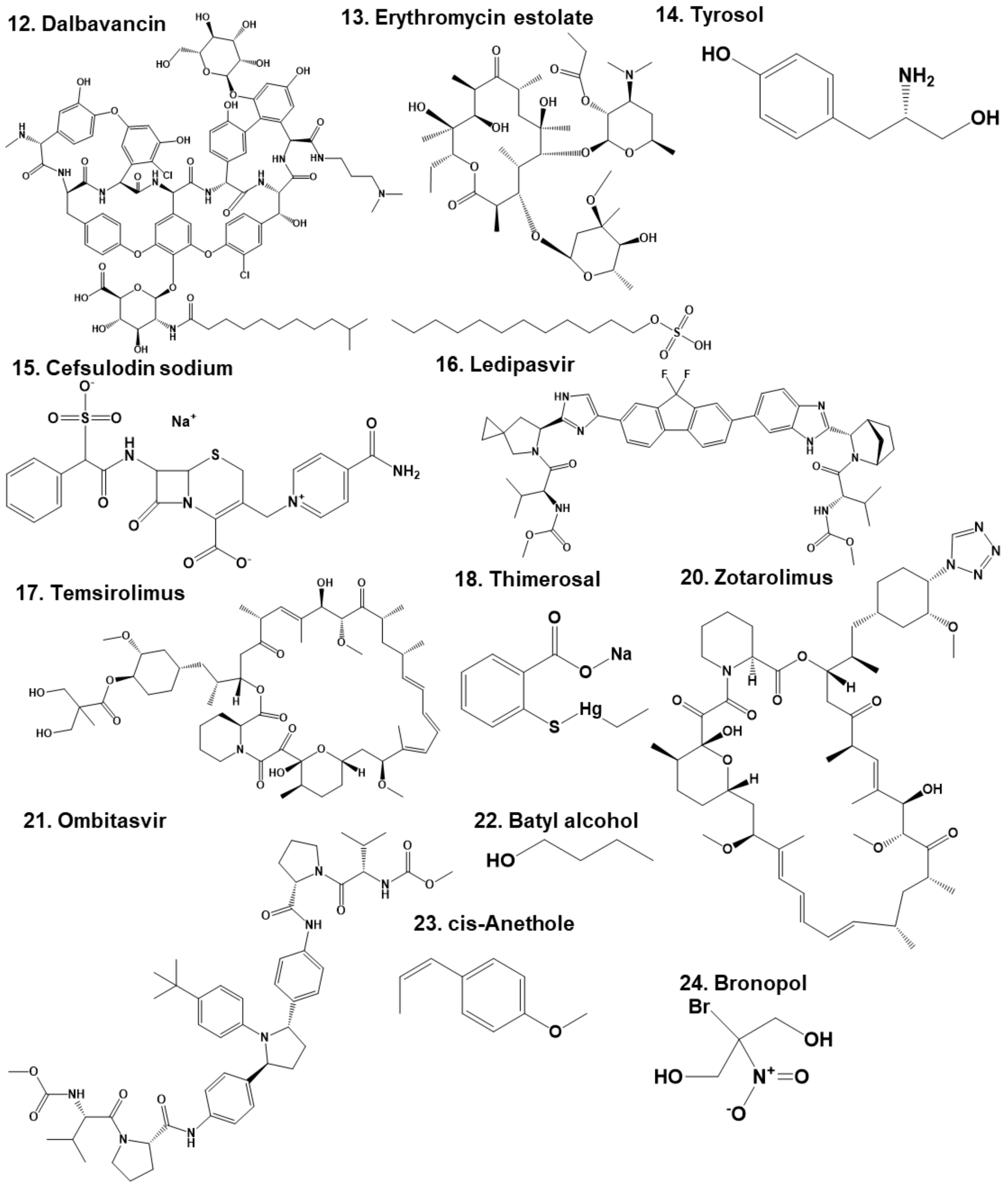


**Figure S4. Chemical structures of Selleck screen Hits.**

**25. Anidulafungin**

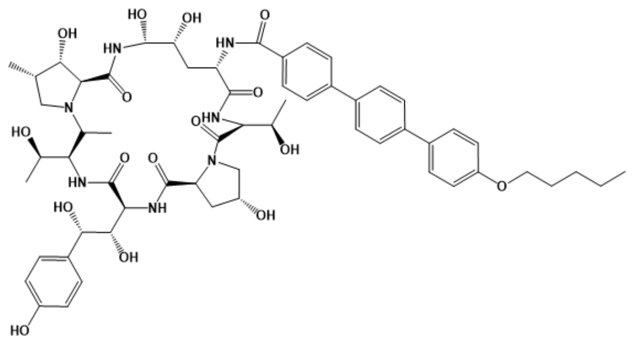

**26. Simvastatin**

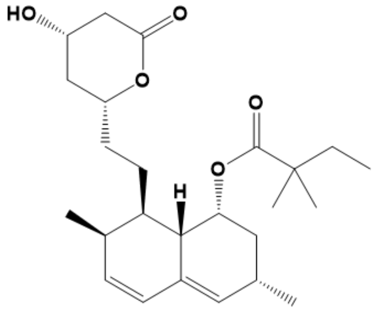

**27. Ridaforolimus**

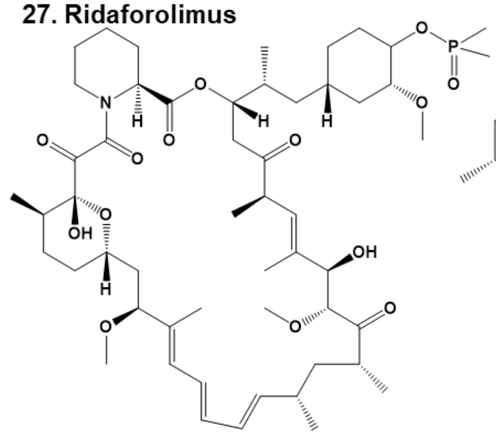

**28. Moxidectin**

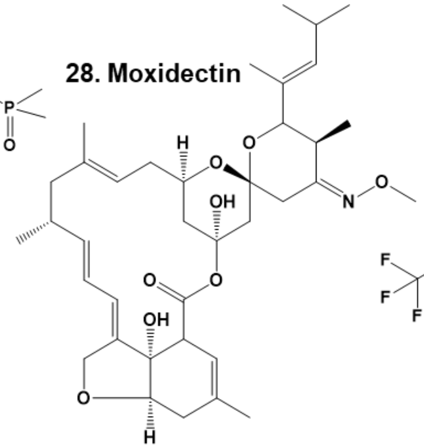

**29. Aprepitant**

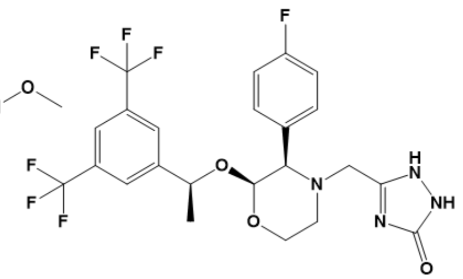

**30. Asunaprevir**

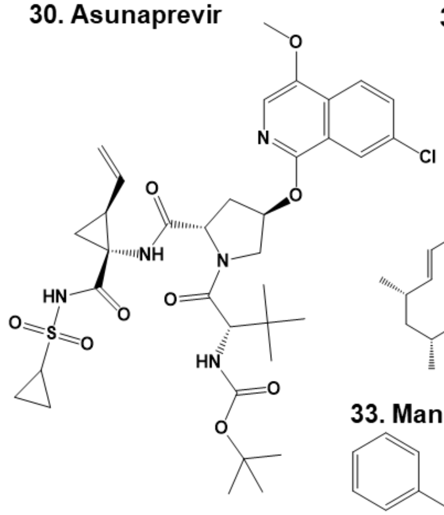

**31. Rapamycin**

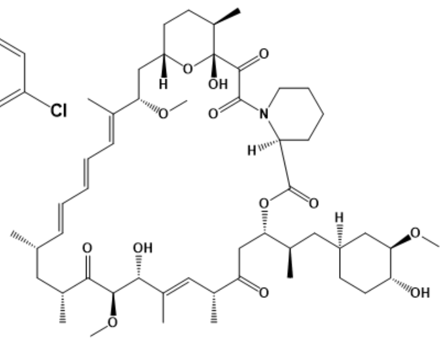

**32. Simeprevir**

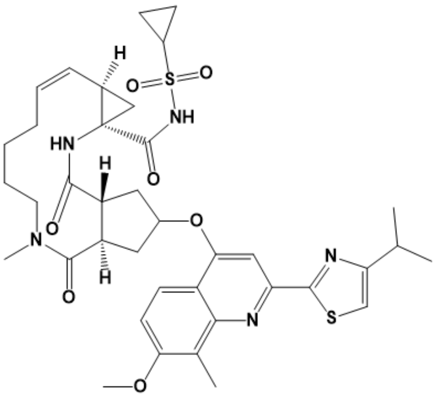

**33. Manidipine**

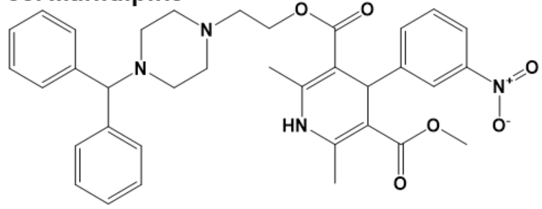

**Figure S5. Chemical structures of Selleck screen Hits.**

**34. Clindamycin palmitate**

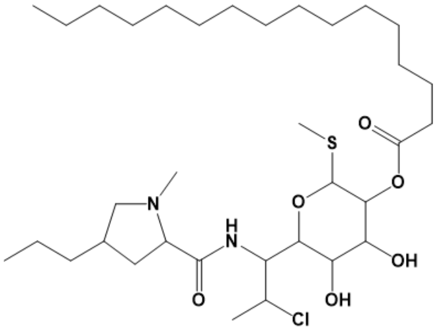

**35. Ertapenem sodium**

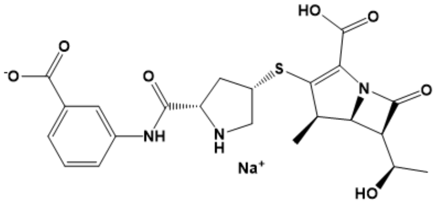

**36. Ethotoin**

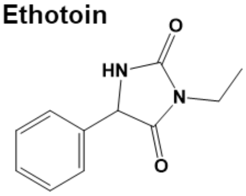

**37. Hydroxyprogesterone caproate**

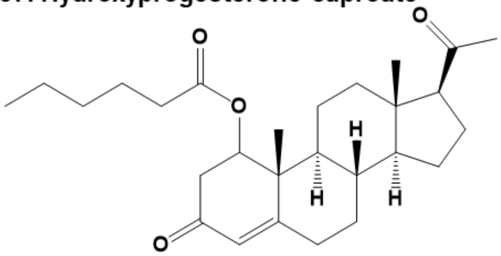

**38. Latanoprost**

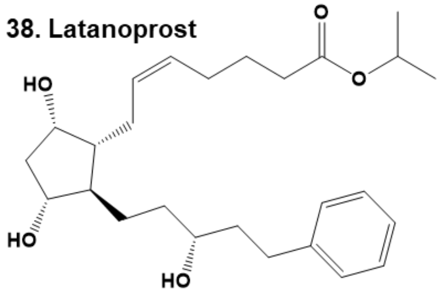

**39. Cilnidipine**

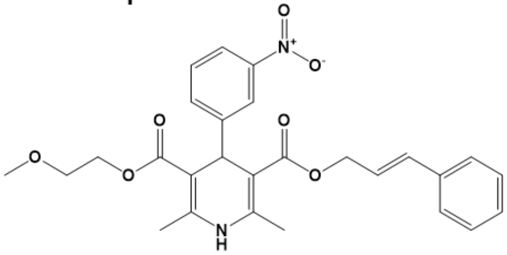

**40. Ouabain**

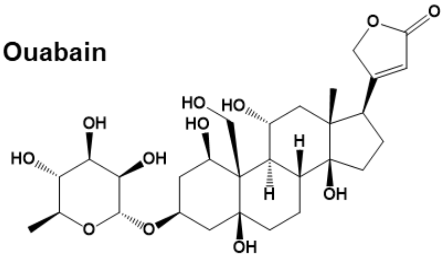

**43. Thiostrepton**

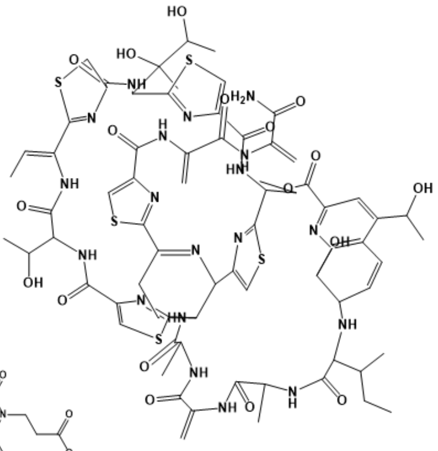

**41. Carfilzomib**

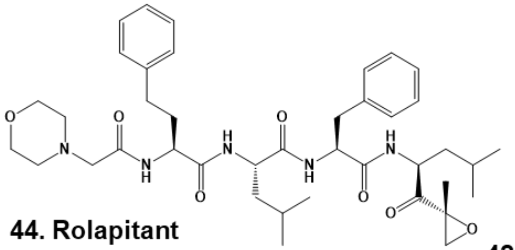

**44. Rolapitant**

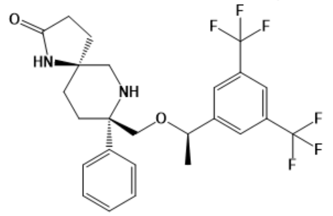

**42. Dabigatran Etexilate**

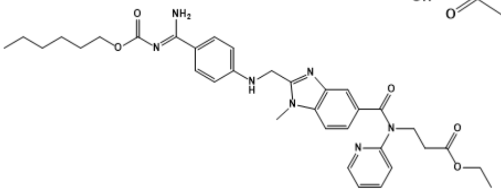

**Figure S6. Chemical structures of Selleck screen Hits.**

**45. Daclatasvir**

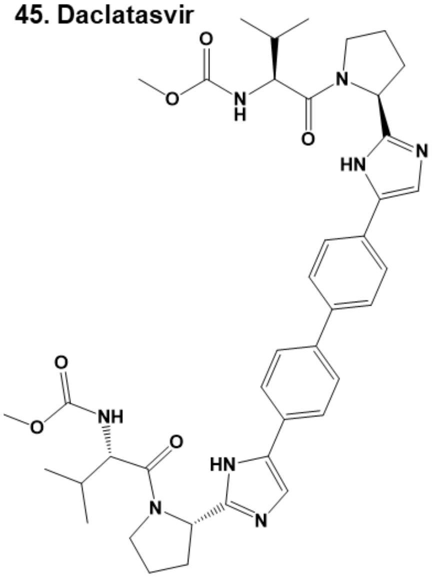

**46. Celecoxib**

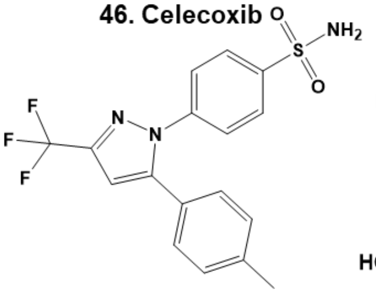

**47. Tafluprost**

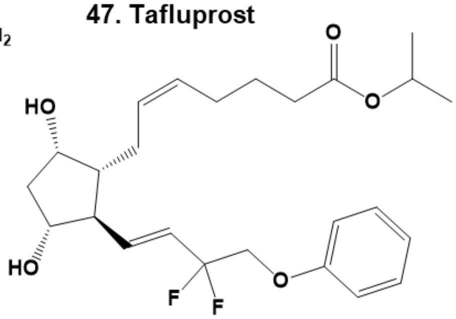

**48. Vorapaxar**

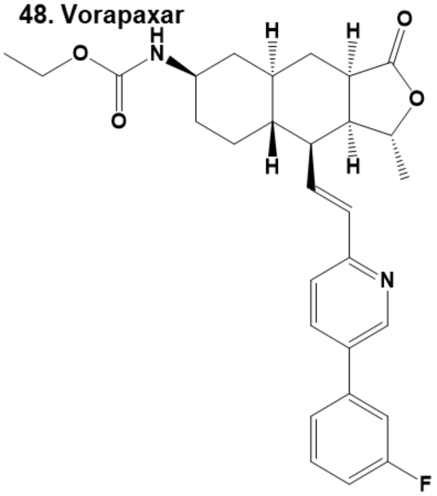

**49. Grazoprevir**

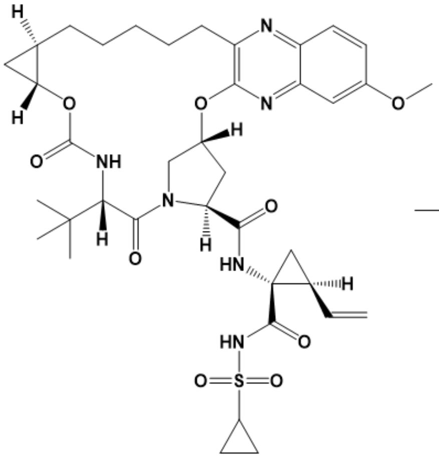

**50. Ibuprofen piconol**

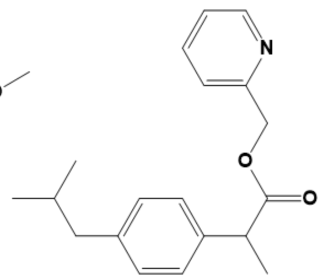

**51. Cabazitaxel**

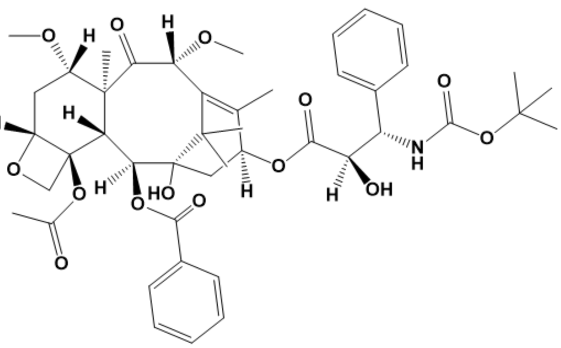

**52. Ivermectin**

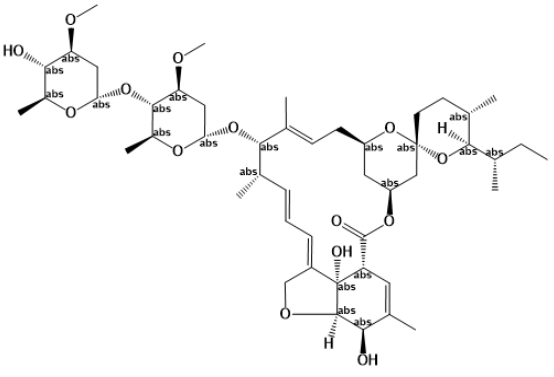

**Figure S7. Chemical structures of Selleck screen Hits.**

**53. Avermectin B1**

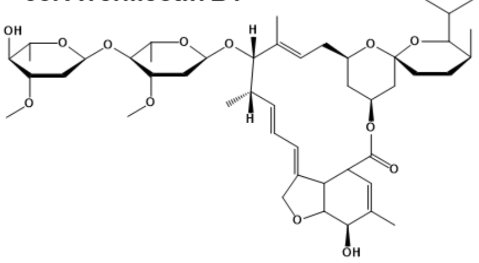

**54. Tacrolimus**

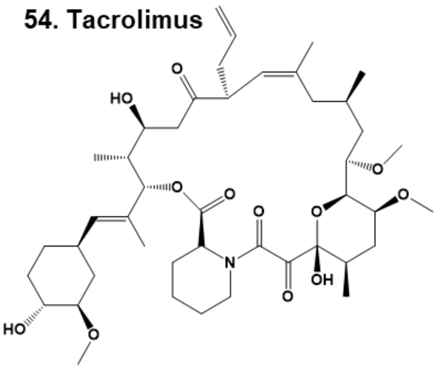

**55. Clonidine**

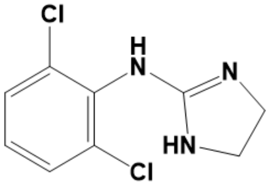

**56. Troglitazone**

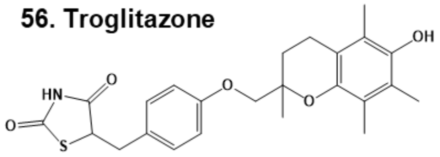

**57. cholecalciferol**

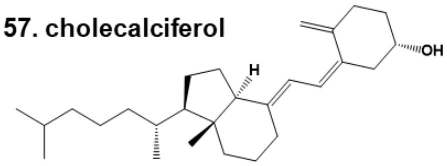

**58. Nefazodone hydrochloride**

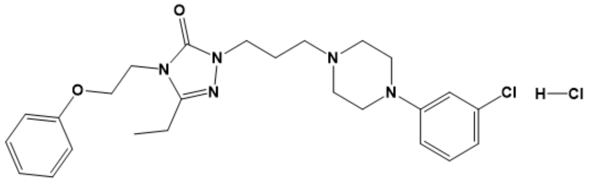

**59. Laurocapram**

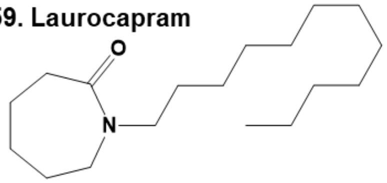

**60. Cabozantinib malate**

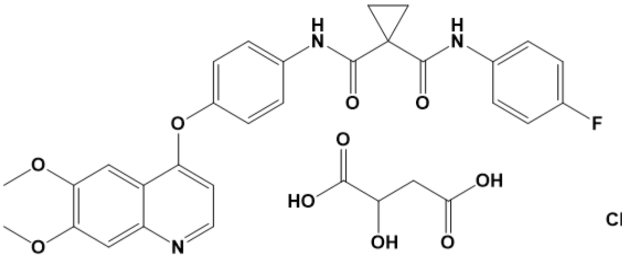

**61. Ketoconazole**

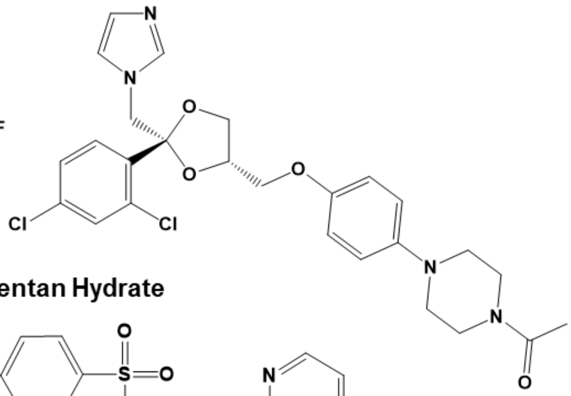

**62. Econazole**

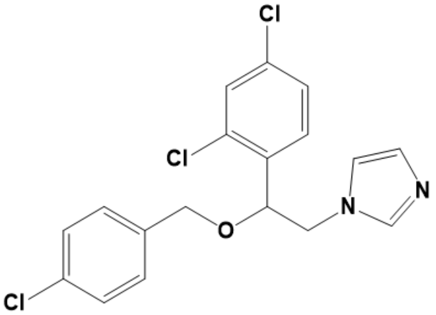

**63. Bosentan Hydrate**

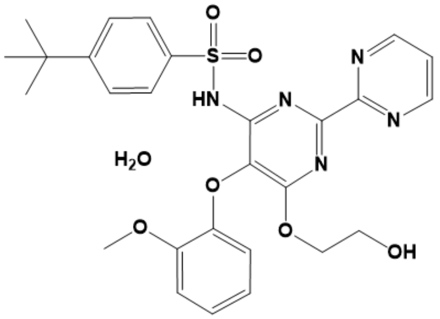

**Figure S8. Chemical structures of Selleck screen Hits.**

64. Avanafil

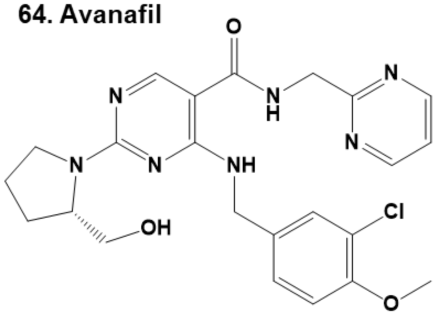

65. Miconazole Nitrate

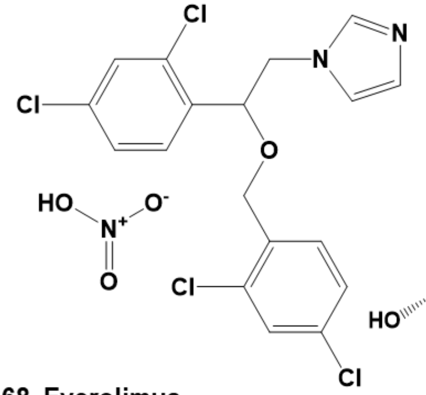

66. Obeticholic Acid

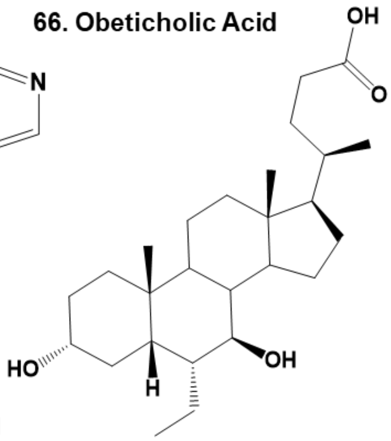

67. Ascomycin

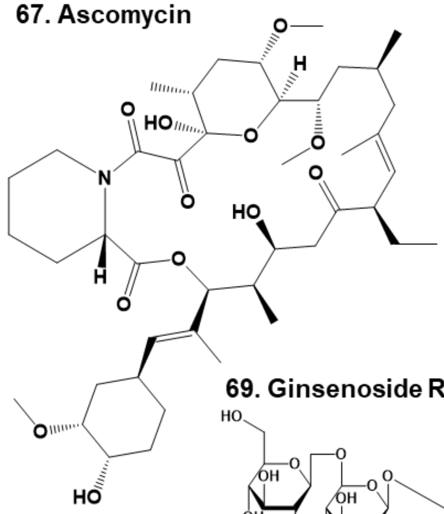

68. Everolimus

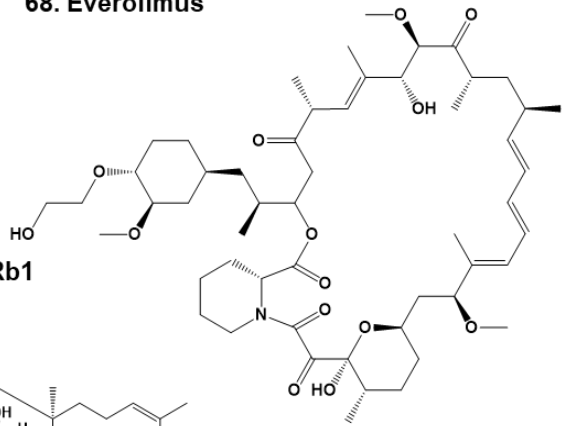

69. Ginsenoside Rb1

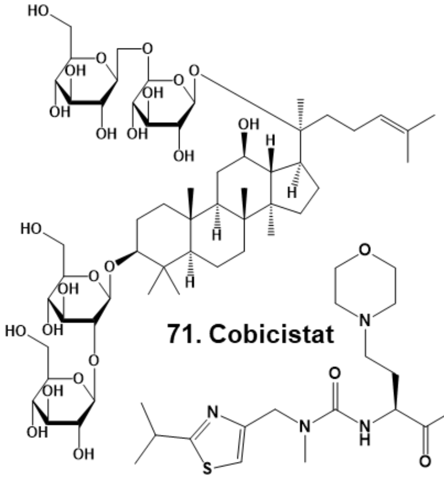

70. Amorolfine HCl

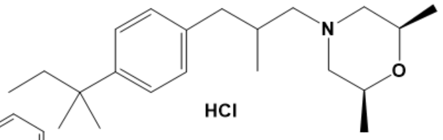

71. Cobicistat

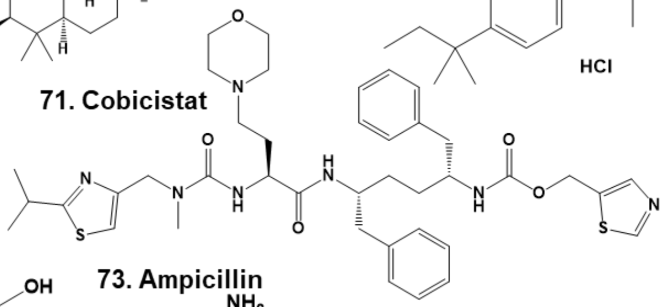

72. Luteolin

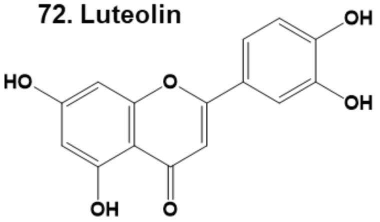

73. Ampicillin

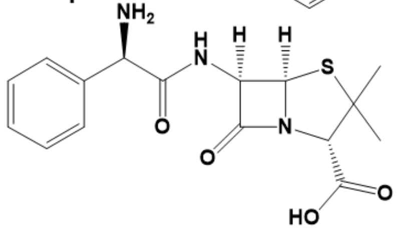

**Figure S9. Chemical structures of Selleck screen Hits.**

74. Nilotinib

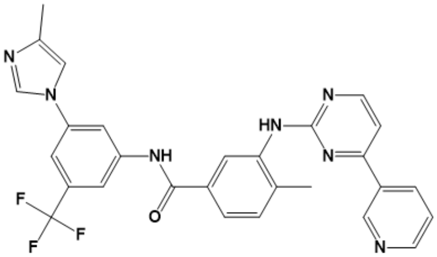

75. Securinine

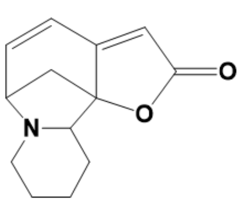

76. Sclareol

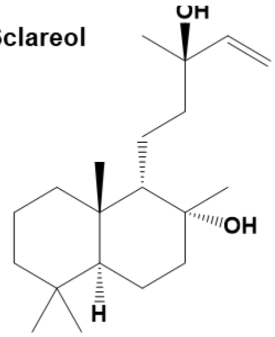

77. Pneumocandin B0

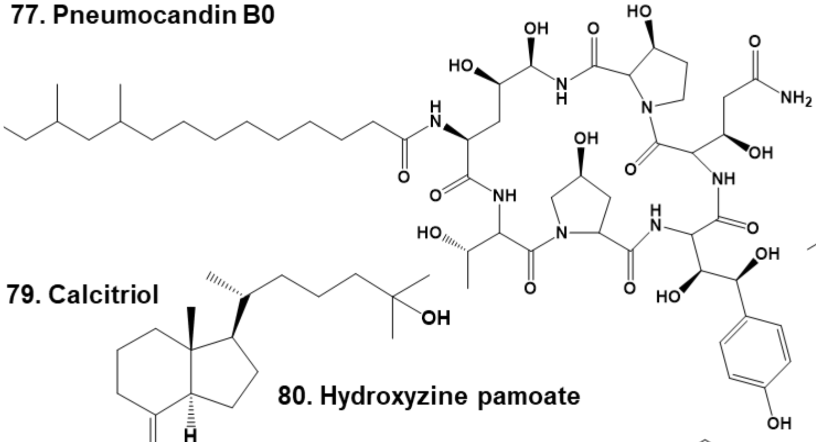

78. AKBA

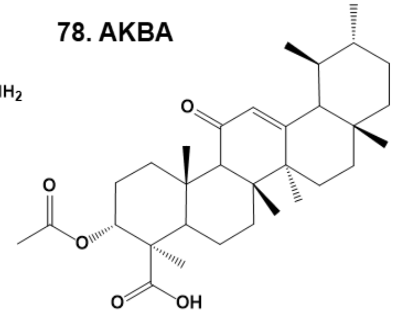

79. Calcitriol

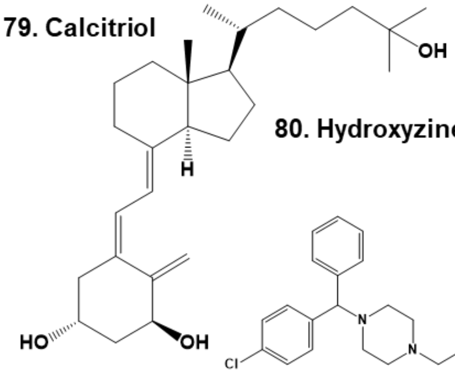

80. Hydroxyzine pamoate

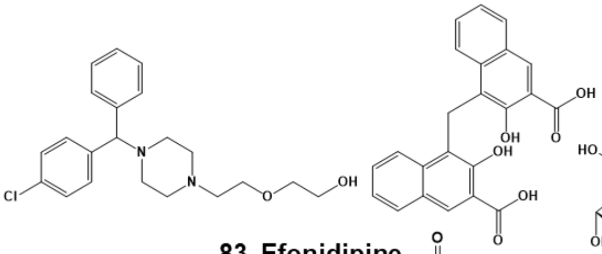

81. Saikosaponin A

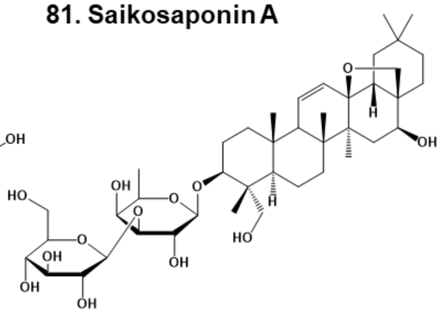

82. Canagliflozin

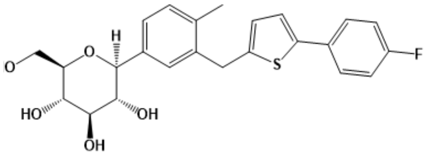

83. Efonidipine

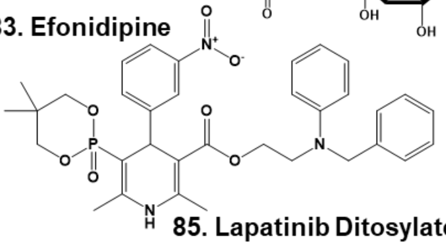

84. Avatrombopag

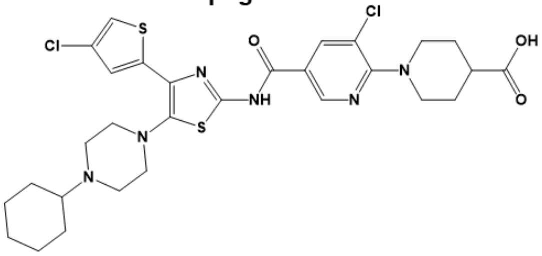

85. Lapatinib Ditosylate

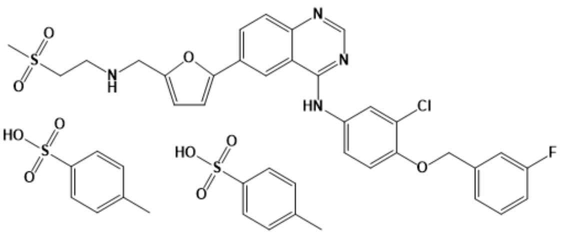

**Figure S10. Chemical structures of Selleck screen Hits.**

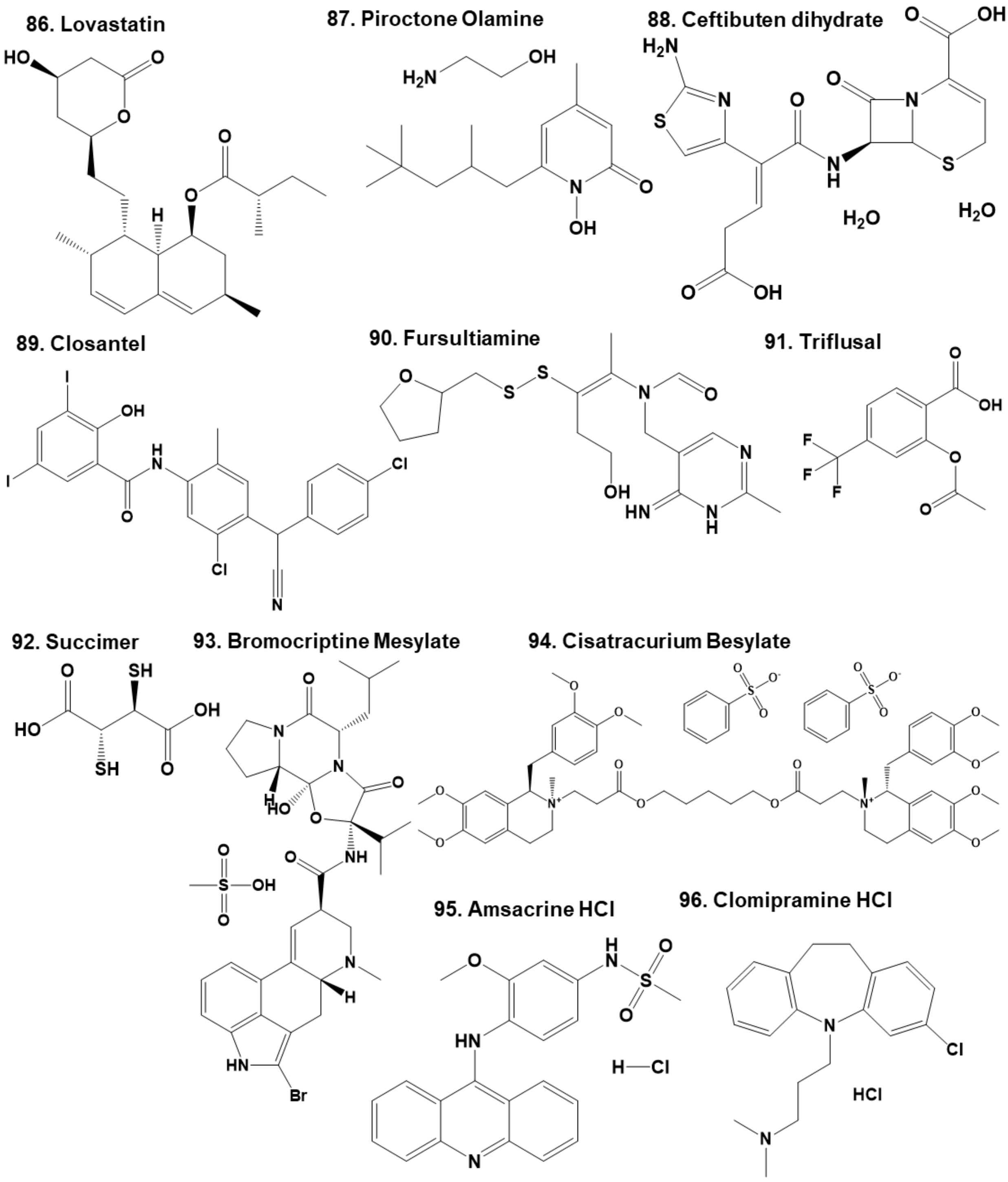


**Figure S11. Chemical structures of Selleck screen Hits.**

97. Primaquine Diphosphate

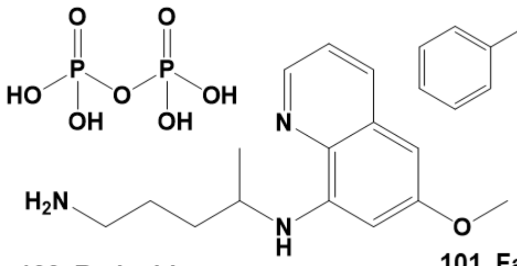

98. Salmeterol

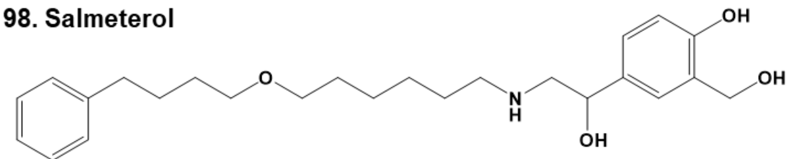

99. Doxycycline Hyclate

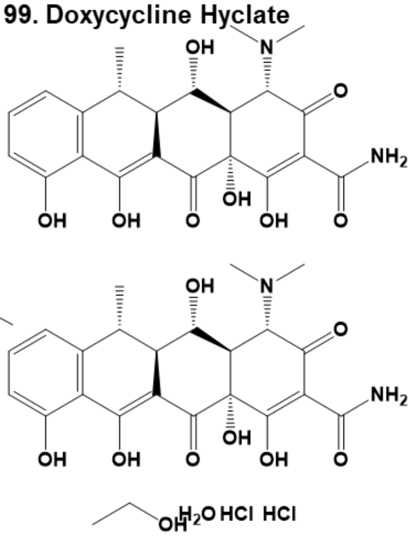

100. Retigabine

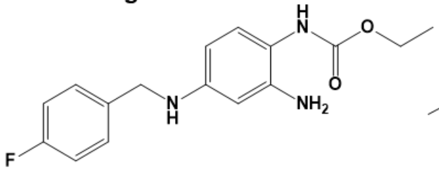

101. Fangchinoline

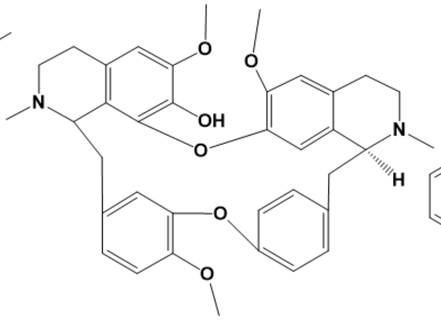

102.Silymarin

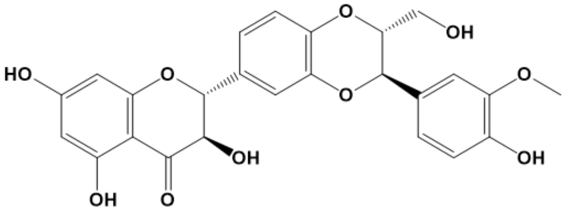

103. Sulbutiamine

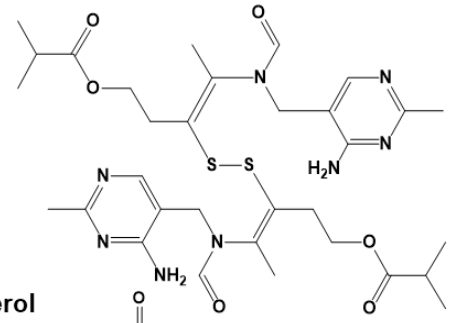

104. Levosimendan

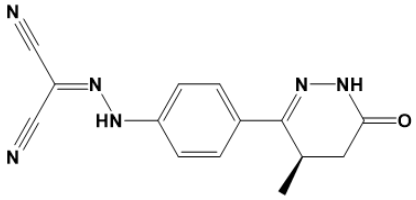

105. Olodaterol

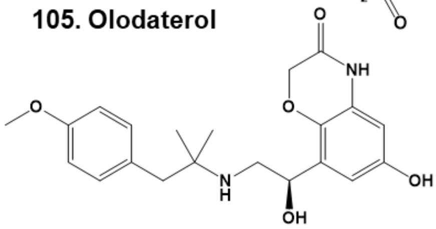

106. Pimavanserin

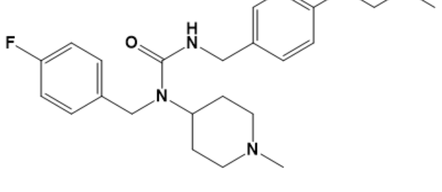

107 and 108. Paroxetine

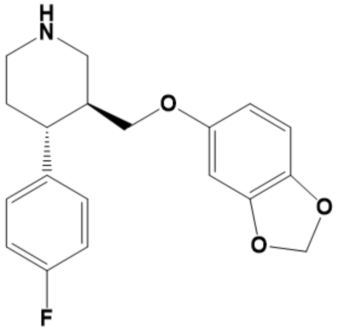

**Figure S12. Chemical structures of Selleck screen Hits.**

109. Nintedanib Ethanesulfonate

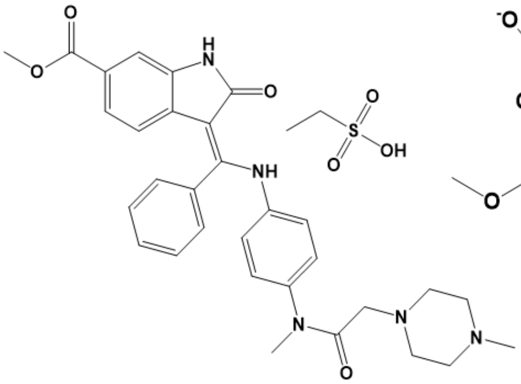

110. Nifedipine

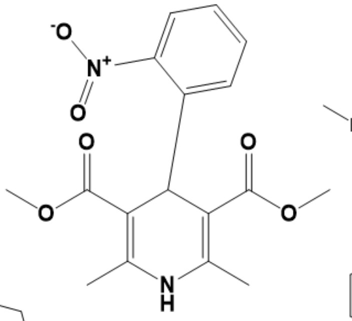

111. pyrvinium

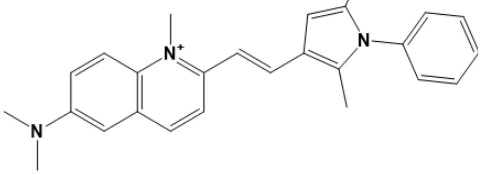

112. Benzalkonium chloride

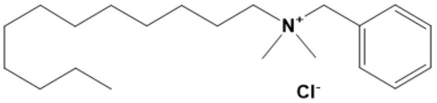

113. Carvedilol

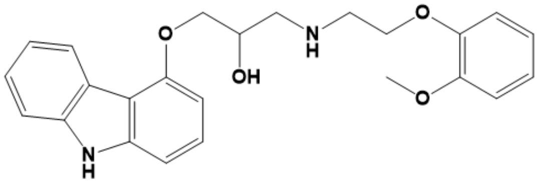

114. (S)-crizotinib

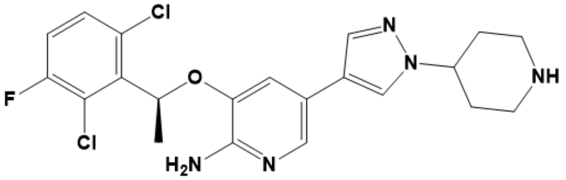

115. Flupirtine maleate

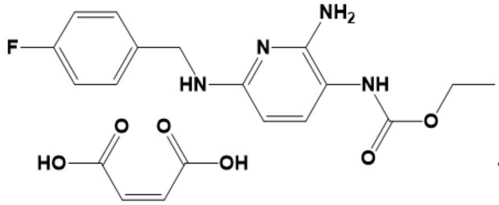

116. Tigecycline

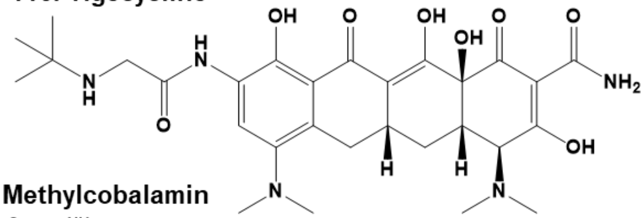

117. Ilaprazole

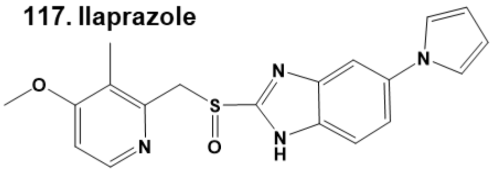

118. Methylcobalamin

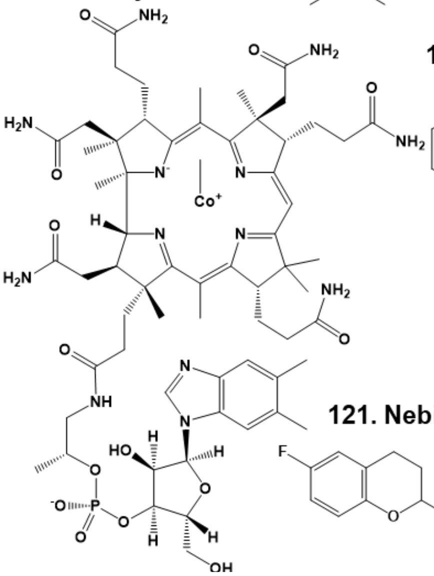

119. CP21R7 (CP21)

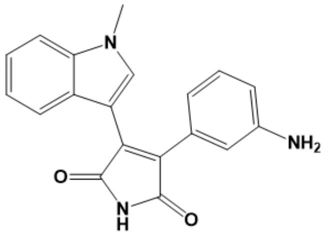

120. Sunitinib Malate

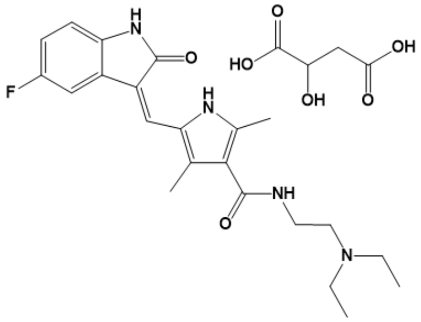

121. Nebivolol

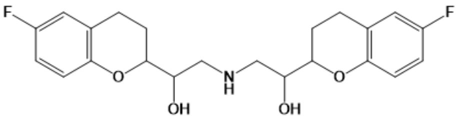

**Figure S13. Chemical structures of Selleck screen Hits.**

**122. (−)-Norepinephrine**

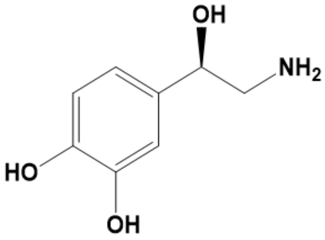

**123. Crizotinib (PF-02341066)**

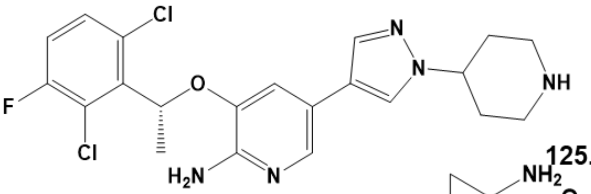

**124. Omadacycline tosylate**

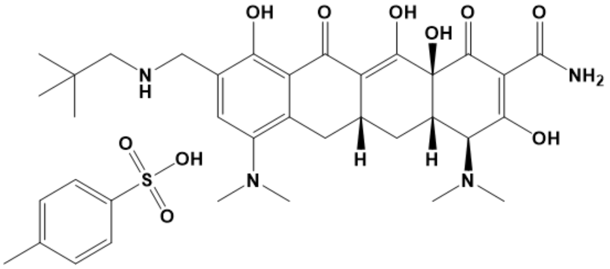

**125. Anlotinib**

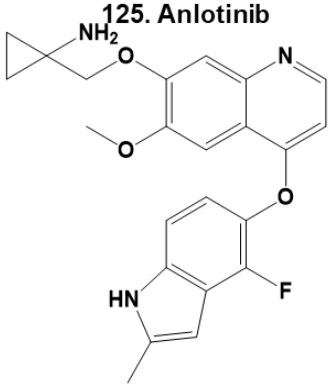

**126. Meclocycline Sulfosalicylate**

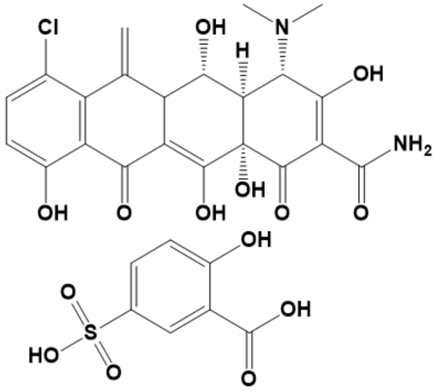

**127. Trichlormethiazide**

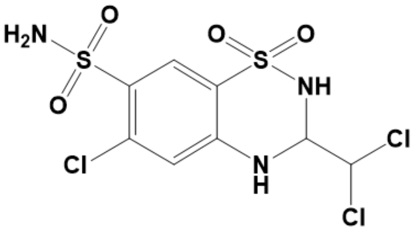

**128. Vortioxetine (Lu AA21004)**

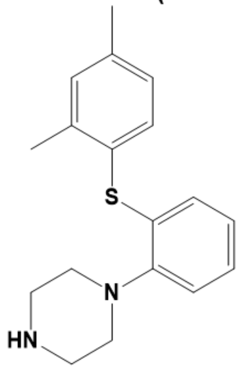

**129. Benzethonium**

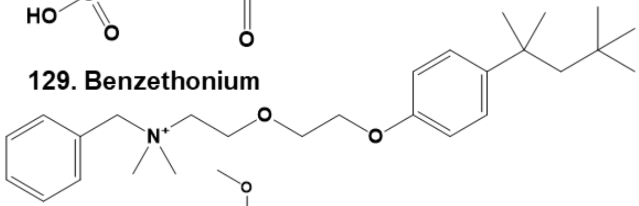

**130. Cyclofenil**

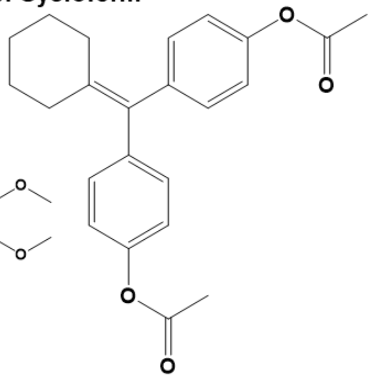

**131. Mivacurium**

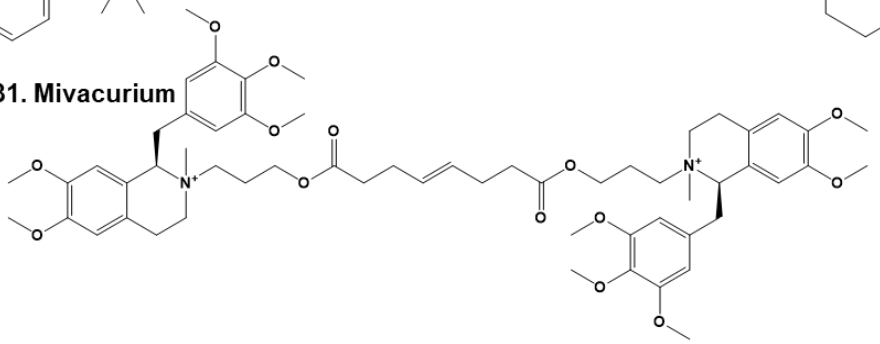

**Figure S14. Chemical structures of Selleck screen Hits.**

**132. Raloxifene**

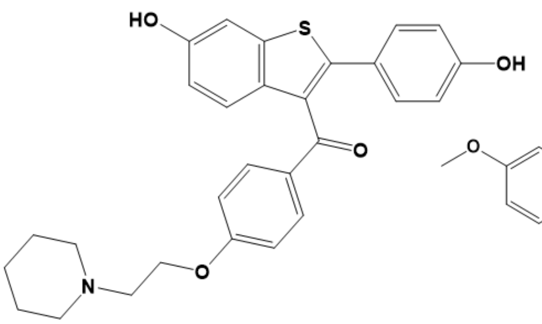

**133. Tegaserod Maleate**

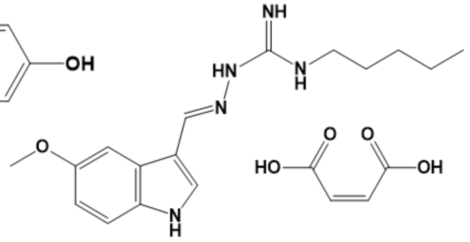

**134. Ethidium bromide**

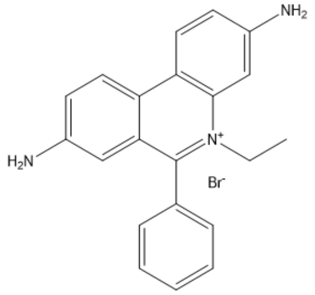

**135. Ethacridine lactate monohydrate**

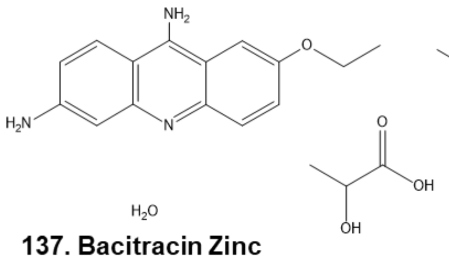

**136. Methylene Blue**

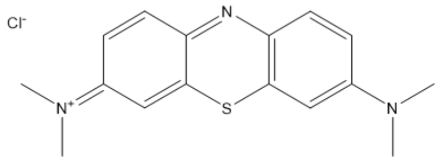

**138. Phenazine methosulfate**

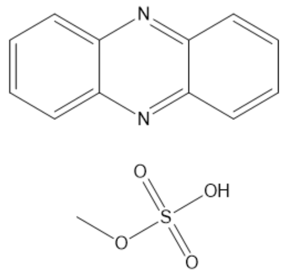

**137. Bacitracin Zinc**

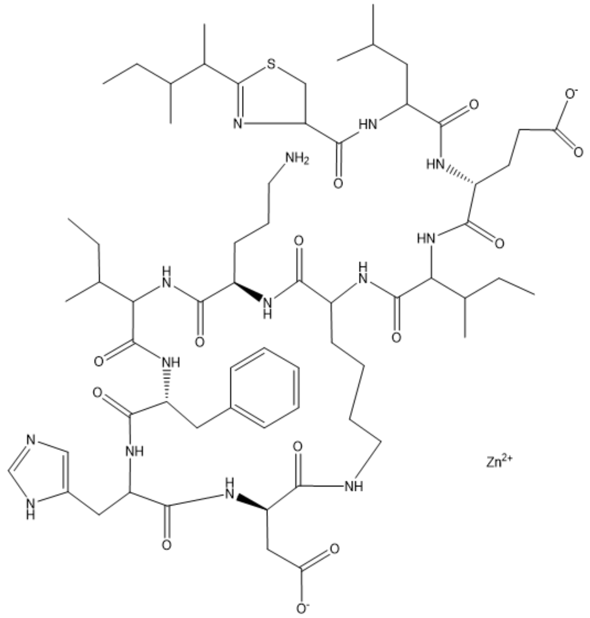

**142. Sanguinarine chloride**

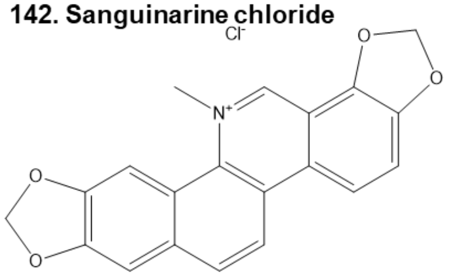

**143. Fingolimod (FTY720) HCl**

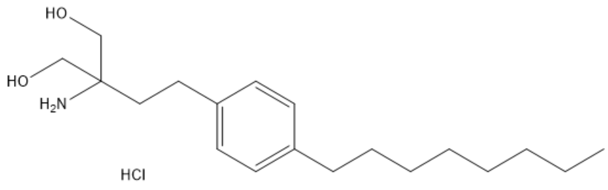

**144. 4-Aminophenol**

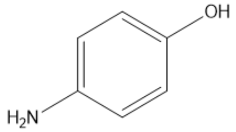

**139 and 141. Chlorhexidine**

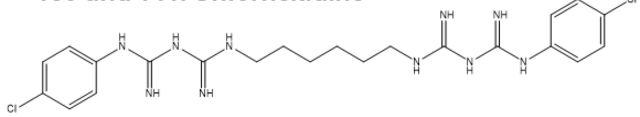

**Figure S15. Chemical structures of Selleck screen Hits.**

**145. Zinc Undecylenate**

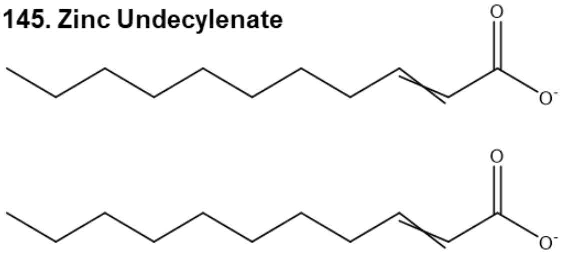

**146. Zinc Pyrithione**

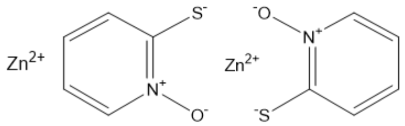

**147. Otilonium Bromide**

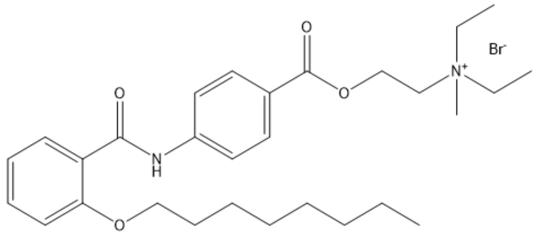

**148. Olanexidine Hydrochloride semihydrate**

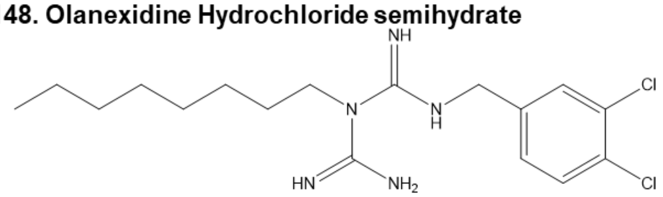

**149. Cetrimonium Bromide**

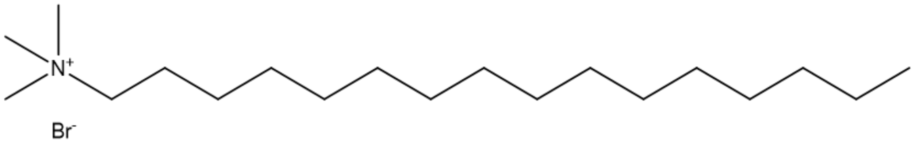

**150. Cetylpyridinium Chloride**

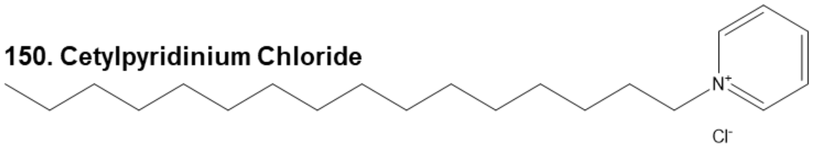

**151. Pixantrone Maleate**

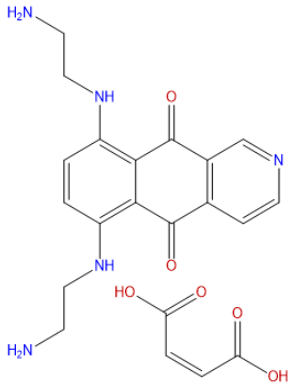

**152. Octenidine Dihydrochloride**

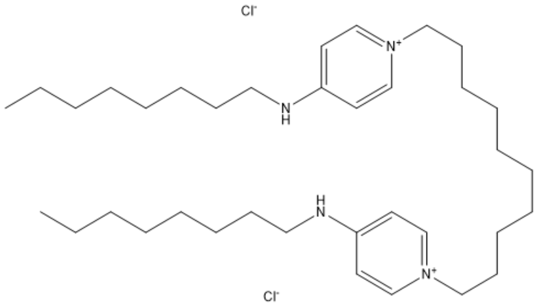

**Figure S16. Chemical structures of Selleck screen Hits.**

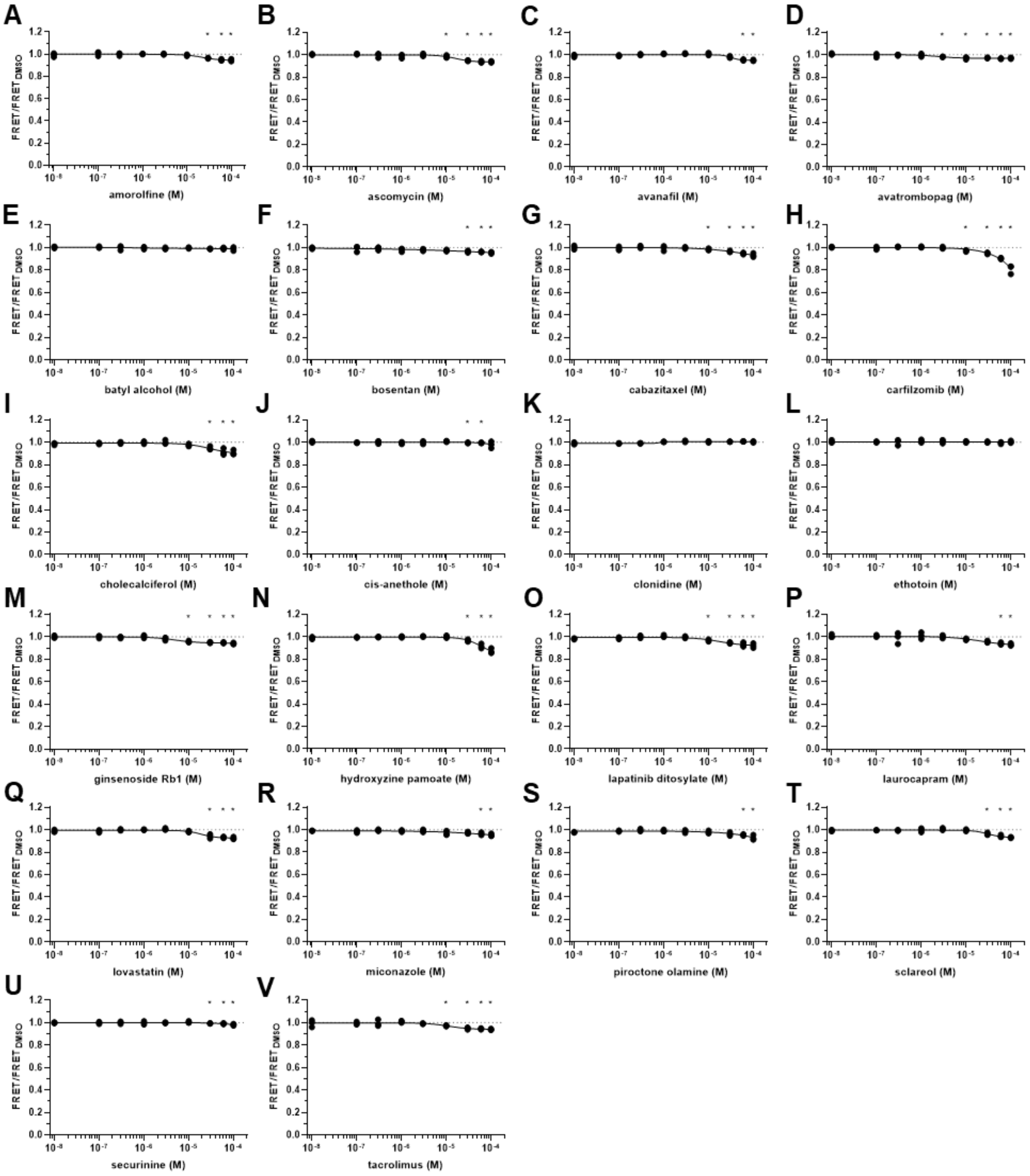


**Figure S17. FRET dose response of Hit compounds that decrease FRET by less than 20%. A-X.** Dose response of Hit compounds amorolfine (**A**), ascomycin (**B**), avanfil (**C**), avatrombopag (**D**), batyl alcholol (**E**), bosentan (**F**), cabazitaxel (**G**), carfilzomib (**H**), cholecalciferol (**I**), cis-anethole (**J**), clonidine (**K**), ethotoin (**L**), ginsenoside Rb1 (**M**), hydroxyzine pamoate (**N**), lapatinib ditosylate (**O**), laurocapram (**P**), lovastatin (**Q**), miconazole (**R**), piroctone olamine (**S**), sclareol (**T**), securinine (**U**) and tacrolimus (**V**) were tested on the in vitro ABD biosensor FRET. Data shown as individual data points, n=3.
